# Uncovering natural variation in root system architecture and growth dynamics using a robotics-assisted phenomics platform

**DOI:** 10.1101/2021.11.13.468476

**Authors:** Therese LaRue, Heike Lindner, Ankit Srinivas, Moises Exposito-Alonso, Guillaume Lobet, José R. Dinneny

## Abstract

The plant kingdom contains a stunning array of complex morphologies easily observed above ground, but largely unexplored below-ground. Understanding the magnitude of diversity in root distribution within the soil, termed root system architecture (RSA), is fundamental to determining how this trait contributes to species adaptation in local environments. Roots are the interface between the soil environment and the shoot system and therefore play a key role in anchorage, resource uptake, and stress resilience. Previously, we presented the GLO-Roots (Growth and Luminescence Observatory for Roots) system to study the RSA of soil-grown *Arabidopsis thaliana* plants from germination to maturity (Rellán-Álvarez et al. 2015). In this study, we present the automation of GLO-Roots using robotics and the development of image analysis pipelines in order to examine the natural variation of RSA in Arabidopsis over time. This dataset describes the developmental dynamics of 93 accessions and reveals highly complex and polygenic RSA traits that show significant correlation with climate variables.

## Introduction

The diversity of shoot and root system forms within the plant kingdom mirrors the many functions required for plant survival in diverse habitats. However, these forms are not static, but dynamically change both over time by evolution and natural selection, but also short term during the lifespan of a plant to acclimate to a changing environment. The visible above-ground shoot is optimized for photosynthesis and reproduction while the below-ground root is responsible for anchoring the plant and allowing it to reach essential nutrients and water. While shoot forms and structures have undergone centuries of scrutiny under the human eye, underground root systems have received much less attention due to impeded accessibility.

Root system architecture (RSA) refers to the spatial arrangement of roots in soil, which encompasses the geometric nature of root connectivity and distribution (Lynch 1995; Rellán-Álvarez, Lobet, and Dinneny 2016). This arrangement of roots depends on relationships between the growth rates, frequency, and orientation of different root types. In allorhizic root systems, common in Eudicotyledons like the model plant *Arabidopsis thaliana*, the embryonically formed primary root is the first to emerge from the seed at germination and ultimately branches to give rise to secondary roots, which, in turn, branch and produce tertiary and higher order roots (Osmont, Sibout, and Hardtke 2007; Fitter 1987). Each step of forming a branch, from the priming of a lateral root primordia to emergence from the parent root, is a tightly controlled process regulated by endogenous and exogenous signals (Moreno-Risueno et al. 2010; Péret et al. 2009; Van Norman et al. 2013; Motte, Vanneste, and Beeckman 2019). Modulation and crosstalk of plant hormones change local growth patterns. The ratios of the phytohormone auxin and its antagonist, cytokinin, control growth throughout the plant and, importantly, regulate primary and lateral root initiation (Tian, De Smet, and Ding 2014; Morris et al. 2017). The hormone abscisic acid (ABA), on the other hand, inhibits lateral root primordia from emerging (De Smet et al. 2003). ABA levels can increase in response to external stress and cause an extended quiescent phase after lateral root emergence (Duan et al. 2013; Xiong et al. 2006). Hence, the precise balance of these hormones influences root development and shapes overall RSA.

In addition to their genetically determined growth program, plants also react to spatial and environmental cues to tune their form for certain environments. Roots must strategically navigate through the soil environment for efficient acquisition of water and nutrients. Understanding the mechanisms by which roots sense and respond to their environment remains a significant challenge since existing methods either compromise on physiological relevance or on throughput, which can impact the ability to deploy functional genomic tools to identify regulatory genes. Most RSA studies are performed on young root systems using easily accessible, *in-vitro* gel-media systems and leverage the powerful genetic and molecular resources of *Arabidopsis thaliana* (Rosas et al. 2013; Ogura et al. 2019; Waidmann et al. 2019; Julkowska et al. 2014). Whereas bigger and more mature root systems of crop species have been studied in soil environments, however, compromising on throughput (Morris et al. 2017; Jiang et al. 2019).

Previously, we developed GLO-Roots (<u>G</u>rowth and <u>L</u>uminescence <u>O</u>bservatory for <u>Roots)</u>, an imaging platform enabling visualization of soil-grown Arabidopsis root systems from germination to maturation by combining custom growth vessels, luminescence-based reporters, an imaging system, and an image analysis suite (Rellán-Álvarez et al. 2015). Here, we present the automation of GLO-Roots by creating GLO-Bot, a robotic platform that enables large scale capture of Arabidopsis root growth over time in a soil-like environment. Along with the automated imaging system, we developed an improved root analysis pipeline for quantification of various RSA traits. Together, these technological advancements enabled time-lapse imaging of RSA development of 93 Arabidopsis accessions from 14 to 28 days after sowing, creating a unique dataset that allows for wide ranging analyses of highly complex, adult root systems. Quantitative genetic analyses and Genome Wide Association Studies (GWAS) yielded insight into the heritability and genetic architecture of distinct RSA traits, and significant associations with climates across the species’ geographic distribution highlighted the power of our approach and its relevance for understanding how root architectural traits contribute to local adaptation.

## Results

### Automation enables high spatio-temporal resolution measurements of root system dynamics

#### Establishing the robotics infrastructure for automated root imaging

In order to further develop the potential of GLO-Roots for time-lapse imaging to track root development from early stages to mature RSAs, we developed GLO-Bot, a cartesian gantry robotic system consisting of an exterior frame and arm on a linear rail system built from T-Slotted aluminum extrusion, which houses the Growth and Luminescence Observatory 1 imaging system (GLO1) ((Rellán-Álvarez et al. 2015) (Figure 1 A, B). In addition to GLO1, the frame contains a growth area for seven black polyethylene growth boxes (Figure 1 B), each of which holds 12 rhizotrons arranged in a two by six grid (Figure 1 C, Supplemental Figure 1 B). In total the system can hold 84 rhizotrons, each one containing one plant.

**Figure 1:**
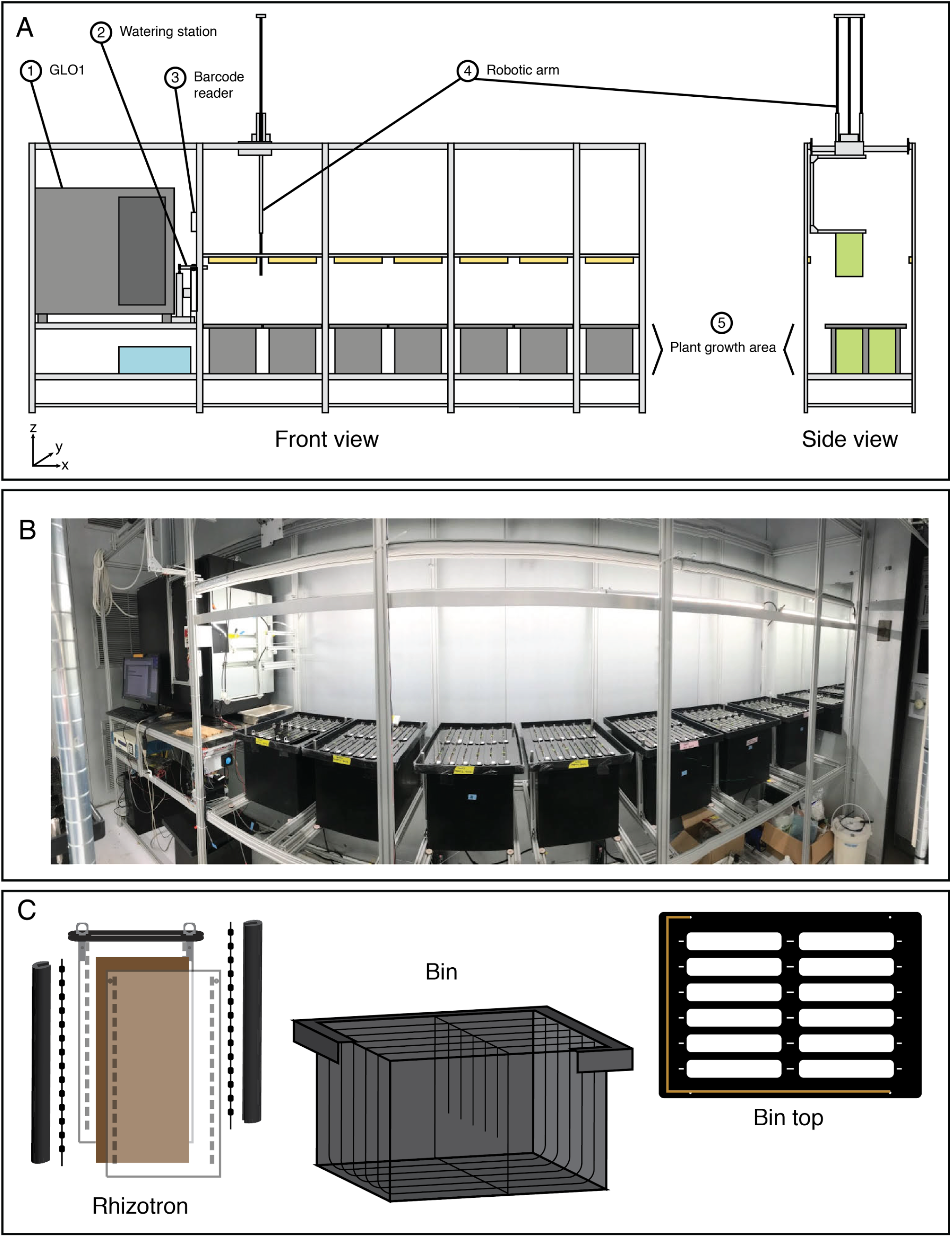
GLO-Roots automation: GLO-Bot. (A) Schematic of the GLO-Bot cartesian gantry system includes: (1) The GLO1 imaging system published in Rellán-Álvarez et al., (2015), which houses cameras and a rotating stage for root imaging, (2) a watering station for general watering and watering with dilute luciferin solution prior to imaging, (3) a barcode scanner to identify the rhizotron and load a specific watering and imaging protocol, (4) a robotic arm, which moves in the x-, y-, and z-directions and has a hook at the end to pick up rhizotrons, and (5) an area for plant growth, which can be seen in the photograph of GLO-Bot (B). (C) Automation updates required updating the GLO-Roots growth vessel designs to include a black acrylic plate and hooks for rhizotron handling as well as dividers within the bins and guides along the bin top, which allow the rhizotron to hang and shield the roots from light. Copper tape along the edge of the bin top enables positioning.

The primary goal of developing the robotics platform was to automate plant imaging since the luciferin watering regime of each rhizotron and the long exposure time required to capture luminescence signal were the most time consuming steps of the GLO-Roots system. An important consideration was the layout for the robot and range of motion for the arm (Figure 1 A). The perpendicular position of the cameras relative to the rhizotron being imaged in GLO1 lent itself well to the final linear rail system and arm with x, y, and z motors. To minimize positional errors associated with finding a particular rhizotron, the robotic arm checks the coordinates of each growth container prior to imaging (Supplemental Video 1). The arm moves to each plant position and lifts the rhizotron via a metal hook, which gives the arm a robust surface with which to pick up the rhizotron (Figure 1 C, Supplemental Figure 1). Both rhizotron and growth box designs were adapted for robotic use in a way that prevents light exposure of the root system during growth (Shi et al. 2018) while also facilitating easy removal and repositioning of rhizotrons in the growth container during imaging. Each rhizotron is shielded from light exposure by creating individual chambers within the growing box (Figure 1 C, Supplemental Figure 1 B). To facilitate removal and replacement of each rhizotron for imaging, we changed the design of the growth box from a standing to a hanging position by adding a piece of acrylic at the top of each rhizotron, which, with the help of small guide pegs, allows the rhizotron to hang in its proper position and shields light from entering the growth container (Figure 1 C).

Programed movements of the robotic arm bring the rhizotron to a barcode scanner, which allows the system to recall the specific treatment and imaging protocol to use (Supplemental Video 1). Subsequently, the arm brings the rhizotron to a watering station where a luciferin solution is applied prior to imaging, which provides the substrate for the constitutively expressed luciferase (Rellán-Álvarez et al. 2015) (Figure 1 A-2). Slow back and forth movements of the rhizotron under a pipette tip connected to a peristaltic pump provides the solution (Rellán-Álvarez et al. 2015) (Supplemental Video 1). As the arm moves, it bypasses a specified gap in the center of the rhizotron to avoid contact with the shoot. The entire watering process takes approximately ten minutes, which allows the water to slowly seep through and spread throughout the rhizotron. The automation was set up such that while one plant is being imaged, the next plant is being watered. To minimize arm movement time, watered rhizotrons are placed on a ledge while the arm removes the imaged rhizotron from the imager and returns it to its growth box (Supplemental Video 1). If the system is running at maximum capacity, GLO-Bot can process 96 rhizotrons in 24 hours, by growing an eighth box on a nearby shelf and swapping out one of the seven boxes in the system during the day.

**Figure 2:**
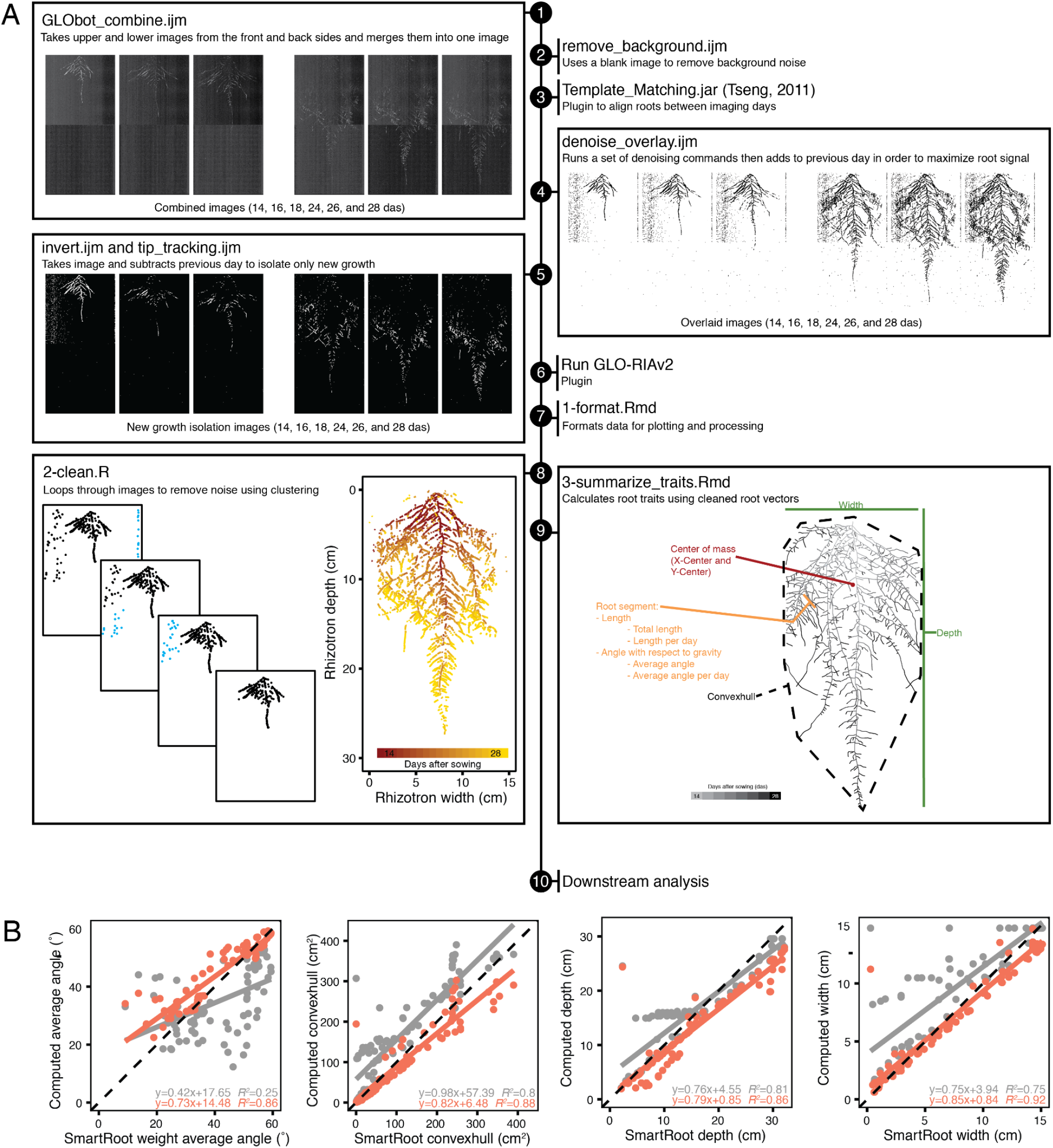
Workflow of image analysis pipeline that enables robust trait measurement. (A) Sample workflow for processing time-series images starts by (1) combining raw images to merge front, back, upper, and lower images, (2) a blank image is then subtracted to remove inherent noise, (3) images are registered to account for small x- and y-movement caused by slight position changes during imaging, (4) registered images are then de-noised and overlaid, which helps overcome luciferase signal loss in older root tissues; however, this step also compounds any background noise. (5) Image subtraction between each day removes this noise and isolates new growth. Processed images are run through GLO-RIAv2 (6) and the output is formatted, such that *in silico* vectorized roots can be reconstructed (7). Additional noise is cleaned up (8) using an iterative distance-clustering method. Traits are then extracted from cleaned roots (9) and the outputs can be used for downstream analysis (10). (B) Comparison of previous root analysis methods (GLO-RIAv1, grey) and the new analysis methods (GLO-RIAv2, coral) with manually traced ground truth measurements (x-axis) reveals that the new analysis method increases accuracy, as demonstrated by the r^2^ values, higher slopes, and lower y-intercepts.

### Development of an image analysis pipeline for dynamic trait quantification

#### Image alignment, signal optimization, and background noise removal

The increased throughput of GLO-Bot enabled time-lapse imaging of root systems and created new opportunities for the measurement of dynamic root growth traits, but also revealed new challenges for root image analysis. Luciferase signal intensity varies between transgenic LUC-expressing lines and decreases in older root tissues, making it ever more difficult to distinguish between roots and noise during the course of the experiment (Figure 2 A-1). While this signal variation is problematic for analysis of raw images, we leveraged our time-lapse data to capture a strong signal for the entire root system by sequentially adding images together. However, in addition to summing the root signal, this step also sums background noise (Figure 2 A-3). Therefore, we utilized a combination of pre- and post-measurement processing to amplify true signal and reduce noise (Figure 2). To account for some of the inherent noise introduced by the imaging system itself, we run the background removal macro remove_background.ijm, to subtract a blank image, taken in GLO1 without a rhizotron present, from every root image (Figure 2 A-2). True root signal can be amplified, or inferred from previous days in the time-lapse, if all images of a root system have the root in the exact same position relative to the exterior image dimensions. Therefore, each series of root images is manually checked to ensure root position remains stationary over time. Misalignments were fixed using the ImageJ plugin Template Matching and Slice Alignment (Template_Matching.jar, (Tseng et al. 2011)). This plugin works by user selection of a region present in all images, a step that might require subsetting of the images, followed by automated detection and translation of subsequent images. During this process, all root images were screened and roots were checked for: (1) appropriate number of images; (2) erroneous image dates; (3) excessive background noise; (4) any other anomalies. Images with excessive background noise, or with too many images, were removed. Next, aligned and cleaned-up images were run through the macro denoise_overlay.ijm, designed to strengthen signal intensity and decrease noise (see detailed steps in material and methods). The output from this macro is a set of images in which each day of imaging is added to the previous days, thus accumulating the strongest signal throughout the root system (Figure 2 A-4).

**Figure 3:**
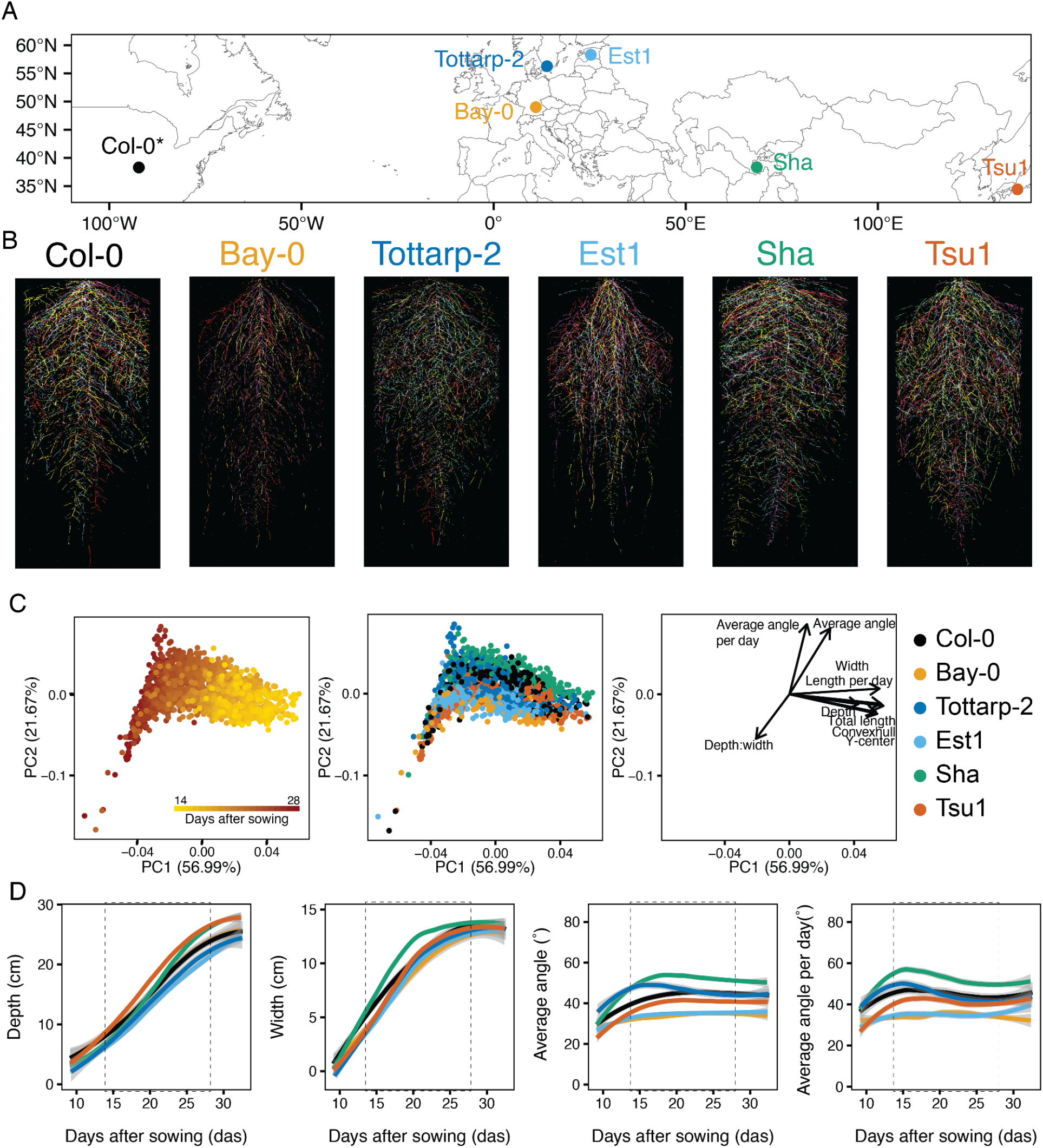
RSA of six Arabidopsis accessions over time. (A) Six accessions from diverse locations were imaged continuously from 9 to 31 days after sowing (DAS). *Col-0 location reflects where accession was originated, not collected. (B) Each accession had a unique RSA pattern (five root systems per accession at 21 DAS each shown in a different color overlaid on top of each other). (C) PCA analysis using eight time points and nine traits shows time (red is 9 DAS and yellow is 31 DAS), as the largest source of variation (PC1) and angle and depth to width ratios further distinguishing the root systems (PC2). When colored by accession, this analysis shows separation between the accessions. Loading plot illustrates the impact of each trait on the overall variation. (D) Four root system traits for six accessions growing over time from 9 to 31 DAS. Trends were calculated using Loess smoothing (n = 12-15 plants at each time point). 95% confidence interval shown by grey shading. Two vertical dashed lines highlight the interval (14 to 28 DAS) chosen for future experiments. Depth and width measurements are constrained by the rhizotron size. Colors correspond to accessions in panel B.

**Figure 4:**
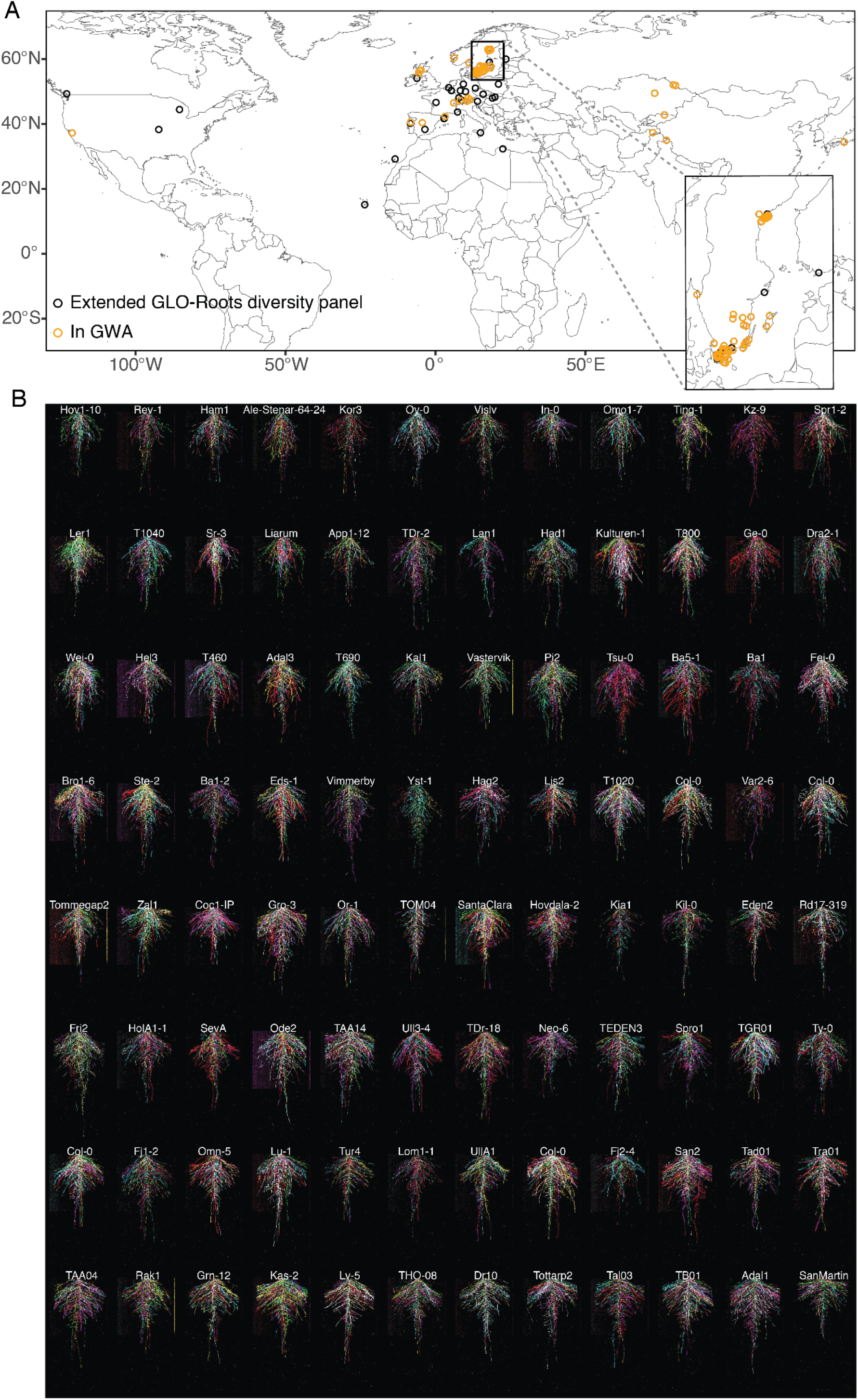
RSA of the GLO-Roots diversity panel. (A) Locations of all accessions used in this study. Inset highlights the concentration of accessions from the Swedish population. (B) Root system architectures of the GLO-Roots diversity panel used in the GWAS at 20 days after sowing (DAS). Six root systems overlaid on top of each other with root system color indicating each replicate. Root systems are arranged in order of median average angle of the root system from deepest in the upper left to shallowest in the bottom right.

With luminescent root signal maximized throughout all of the images in a time-lapse series, the macro invert.ijm is used to invert the images for whole root system analysis. In addition to determining architectural parameters of complete root systems, we also established methods to quantify the actively growing regions of root systems. For this, we use the macro tip_tracking.ijm to isolate the new growth between images by a series of dilating and subtracting of two consecutive images (see detailed steps in material and methods; Figure 2 A-5).

**Figure 5:**
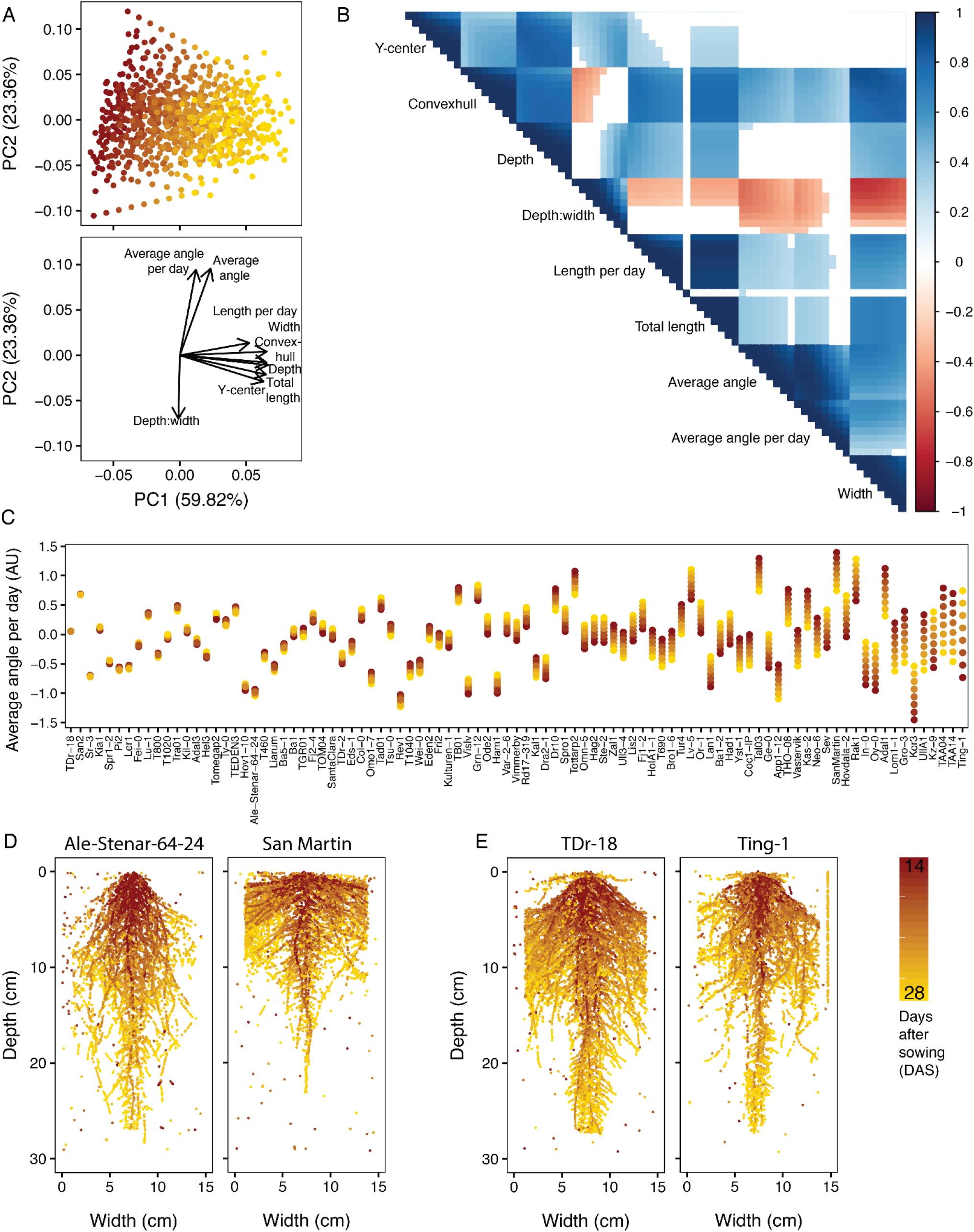
Trait relationships amongst the GLO-Roots diversity panel. (A) PCA analysis of nine fitted traits for the 93 accessions of the GLO-Roots diversity panel reveals traits influenced by time as the predominant source of variation within the dataset (PC1), while angle measurements and the depth to width ratio further distinguish accessions (PC2). (B) A correlogram of the calculated trait values at each time point, increasing left to right and top to bottom, further shows the negative correlations between the depth to width ratio and the angle traits and displays the relationships between all nine traits at each time point. p < 0.01 for all correlations displayed. While some traits remain stable over time, other traits change over time. The extent to which a trait changes varies between accessions. (C) In some accessions, the angle of new growth (average angle per day) remains constant each day, while in others the angle of new growth increases or decreases over time. (D) Ale-Stenar-64-24 and San Martin demonstrate contrasting root architectures, with opposite depth to width ratios and opposite average angles. (E) TDr-18 and Ting-1, on the other hand, are examples of accessions with constant and variable (respectively) average angle per day measurements.

#### Downstream analyses of root growth dynamics requires the distinction between true root signal and background pixels

All images were run through a modified version of GLO-RIA (Growth and Luminescence Observatory Root Image Analysis) ((Rellán-Álvarez et al. 2015), GLO-RIAv2 (Figure 2 A-6). GLO-RIAv2 removes features deprecated from the 2017 ImageJ update (Schneider, Rasband, and Eliceiri 2012; Schindelin et al. 2012) and includes root angle output ranging from 0-180°.

**Figure 6:**
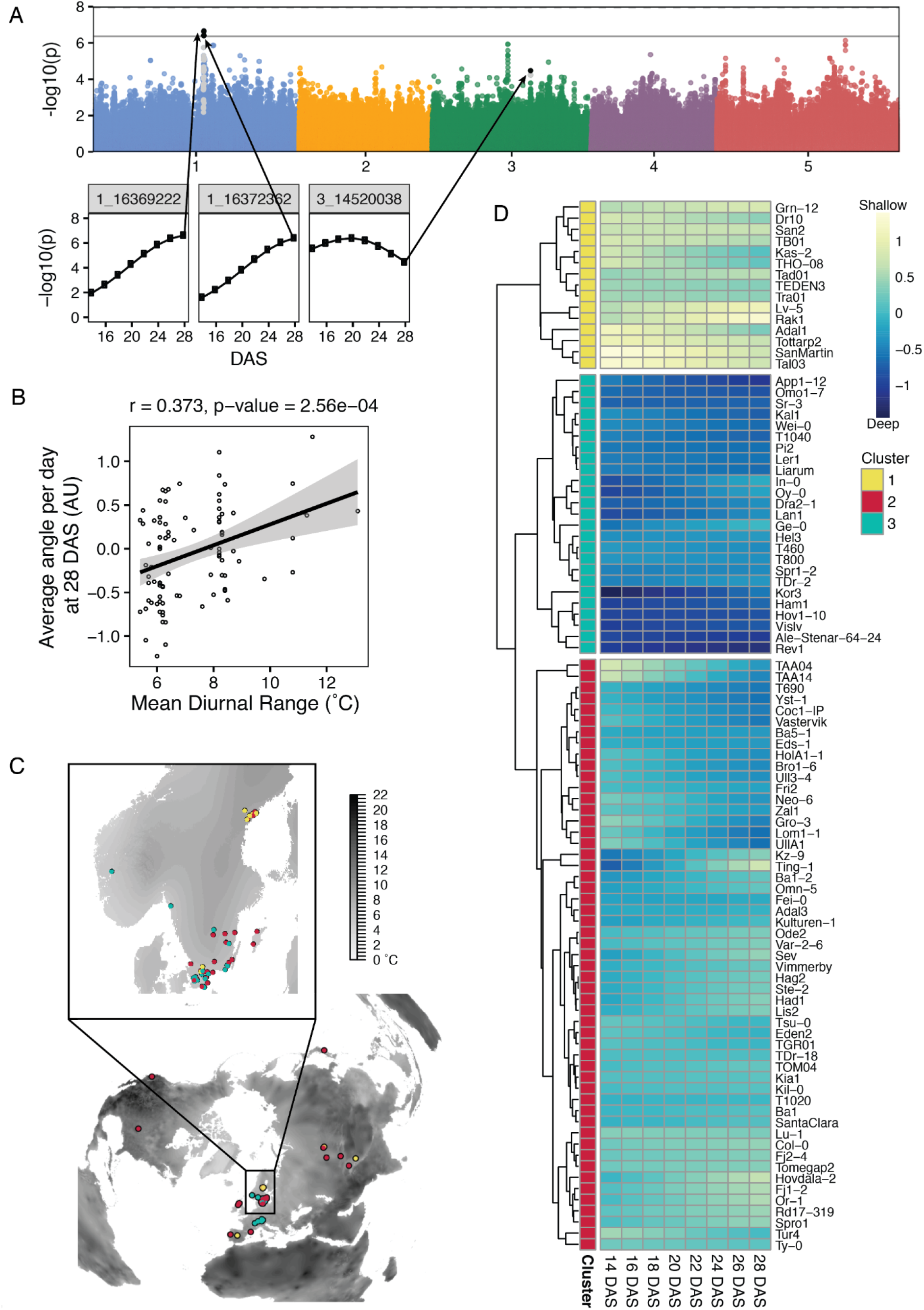
GWAS and Climate correlations of root traits of the GLO-Roots diversity panel. (A) Manhattan plot for average angle per day at 28 days after sowing (DAS). Black points indicate SNP positions that pass the Bonferroni threshold (solid line) at least once throughout the time series, as shown in the inset plots. Grey points are those in linkage disequilibrium (LD) with the significant SNPs. (B) Correlation between the average angle per day and the mean diurnal temperature range, indicates the potential importance of root angle for surviving highly variable climates. (C) A world map showing mean diurnal range (black represents larger fluctuations) with the distribution of accessions (points) colored by cluster identity based on changes in average angle per day over time and computed by between-group average linkage hierarchical clustering with distance = 1.75, as shown in the heatmap (D). Heatmap depicts average angle per day for each day with blue indicating steeper angles and yellow indicating shallower angles. Changes in average angle per day divided the accessions into three clusters: the yellow cluster with consistently shallow root growth, the blue cluster with consistently deep root growth, and the red cluster with intermediate or changing root angles.

Downstream analyses of the GLO-RIAv2 outputs were performed in R (R Core Team 2019). We first process the GLO-RIAv2 output to give us root segment data including 1) the x-, y-coordinate of the upper left corner of the bounding box for each segment, 2) the segment length, and 3) the angle with respect to gravity, such that root segments can be referenced with 0° pointing straight down and 90° pointing straight out to the side. These data give us the ability to reconstruct the root system *in silico* as a series of vectors (Figure 2 A-7).

Next, we remove non-root particles by iteratively clustering the starting points of the measured segments. Clusters are tested for distance between points, the number of segments, and the linearity of the points within the cluster (detailed steps in materials and methods, Figure 2 A-8). Segments that do not meet empirically defined criteria are removed, leaving behind only “true root” segments. Using the “true root”, we calculate traits associated with new growth including: number, length, and angle of new root segments, as well as traits at the whole root system level: width, depth, center of mass, convex hull, depth to width ratio, total length, and average angle (Figure 2 A-9). Together, the automated image analysis pipeline represents a robust tool for dynamic root trait analysis.

To test the accuracy of our trait extraction methods, we used SmartRoot (Lobet, Pagès, and Draye 2011) to manually trace each root system and ground truth our parameters. We observed a strong correlation between our traits extracted via our new image processing pipeline and those that were measured manually (Figure 2 B). While the initial GLO-Roots pipeline works well for end-point image analysis, the above mentioned updates allow for more accurate root trait measurement of time course images (Figure 2 B) and gives us ways to explore new temporal traits.

#### Time-lapse imaging of six Arabidopsis accessions reveals distinct root growth trajectories

In order to test our automated system and to reveal if we can distinguish genotype-specific growth patterns during RSA development over time, we examined a set of six Arabidopsis accessions based on their geographic dissimilarity and available genetic resources (Figure 3 A). Each accession was grown in ten replicates, half of which were imaged from 9 days after sowing (DAS) until 31 DAS and the other half from 21 – 31 DAS. The application of our newly developed imaging protocols to these data allowed us to optimize experimental parameters, including luciferin watering frequency, imaging frequency, and imaging time period. The six accessions that were imaged displayed diverse RSAs (Figure 3 B, Supplemental Video 2) and principal component analysis (PCA) demonstrated that we could distinguish the accessions based on nine of the traits extracted using our image analysis pipeline (Figure 3 C, Supplemental Table 1). Expectedly, time accounted for the largest source of variation (Figure 3 C), indicating that the developmental stage of a plant is the most important variable affecting RSA. We could see that depth to width ratios and average angle measurements further separated the accessions (Figure 3 C). Looking at the traits over time demonstrated that each of the accessions had its own unique growth trajectory with average angle and average angle per day exhibiting the most diversity (Figure 3 D, Supplemental Figure 2). Early during plant growth all accessions have fairly similar average angles, presumably because they have few lateral roots, but over time their average angles for the whole root system, with respect to gravity, shift and distinguish the accessions. By looking at these data, we established that most traits stabilized by the end of the 28 DAS growth period (Figure 3 D, Supplemental Figure 2). Importantly, having imaged the plants both continuously and only at 21 DAS, we were able to confirm that these traits were not influenced by continuous imaging and the associated luciferin watering (Supplemental Figure 3). Time-lapse imaging of these six accessions proved the robustness of GLO-Bot over several weeks, the ability of our image analysis pipeline to determine various root system traits over time, and that even a small subset of accessions showed substantial variation in RSA traits.

### Natural variation in dynamic root system architecture traits is highly complex and polygenic

#### The GLO-Roots diversity panel

To further leverage natural variation and, thus, gain insights into local adaptation of RSA dynamics, we transformed an initial set of 192 *Arabidopsis thaliana* accessions with the *ProUBQ10:LUC2o* reporter for use with the GLO-Roots system (Rellán-Álvarez et al. 2015) (Supplemental Figure 4 A). These accessions comprise the extended GLO-Roots diversity panel, which originated from a densely sampled Swedish population (Long et al. 2013) and were supplemented with accessions from extreme environments with high and low annual average precipitation (Fick and Hijmans 2017), as well as accessions with high and low sodium accumulation levels in the shoot (Figure 4 A, Supplemental Figure 4 B) (Baxter et al. 2010). Ultimately, the GLO-roots diversity panel, a set of 93 accessions, were selected for the GWA population (Figure 4 A, Supplemental Table 2). This set balances genetic similarity and environmental variation along with reporter expression and the number of independent transformation events recovered. This population not only represents a foundational resource for describing phenotypic trait diversity in the Arabidopsis species, but can later be used to identify genes underlying this trait diversity over time, using the powerful tool of Genome Wide Association mapping.

To capture the breadth of RSA dynamics within the diversity panel, we grew six replicated individuals per ecotype in GLO-Bot and imaged every other day from 14 to 28 DAS (Figure 4B, Supplemental Video 3, Supplemental Figure 4 C). Next, using our image analysis pipeline, we focused on the nine root traits that distinctly separated RSAs in the subset of six accessions, and which encompass RSA at a broad scale (Figure 4 B, Supplemental Table 1). For each of these traits, we used a generalized linear mixed model in the R package MCMCglmm to account for replicate and block effects noises and estimate the “breeding value” of each accession for a given trait (Wilson et al. 2010; Mrode 2014); (Hadfield 2010) (Supplemental Figure 5). The denoised trait values for each genotype at 0, 48, 96, 144, 192, 240, 288, and 336 hours after the start of imaging (equivalent to 14, 16, 18, 20, 22, 24, and 28 DAS) were used for further analyses.

#### Root trait diversity and phenotypic relationships amongst the GLO-Roots diversity panel

Similar to the analysis of the first set of six accessions (Figure 3 C), PCA of the GLO-Roots diversity panel revealed that the first principal component captures trait variation over time (Figure 5 A, Supplemental Figure 6 A), as well as traits that are strongly associated with time and, thus, developmental stage (Figure 5 A, Supplemental Figure 6 B) such as the root system’s area (convex hull area), length (total length), and depth. The second principal component demonstrates that both the average angle of the root system, and the average angle of new growth, account for a large amount of root system variation (Figure 5 A, Supplemental Figure 6 C, D). Average root angle and depth-to-width ratio are negatively correlated (Figure 5 A, B), as demonstrated by the accession San-Martin, which has the smallest depth-to-width ratio and the highest average root angle (i.e. most shallow) as well as Ale-Stenar-64-24, which has one of the highest depth to width ratios and lowest average root angles (Figure 5 D). Additional data exploration and accession comparisons can be done using the R shiny App (https://tslarue.shinyapps.io/rsa-app/), which plots the vectorized root systems for all of the accessions.

Further analysis of the traits via pairwise comparison at all time points not only allowed us to visualize existing correlations within traits over time, but also amongst traits (Figure 5 B). Looking at average angle per day, we see that consecutive days show strong correlation across accessions (e.g. day 14 compared to day 16: Spearman’s rank correlation 0.99), whereas later time points are less well correlated with early time points (day 14 compared to day 28: Spearman’s rank correlation 0.56, Figure 5 B). This suggests that the average angle of lateral root tips is a trait that changes over time with some accessions showing greater change than others (Figure 5 C). Indeed, examining this trait across our 93 accessions reveals great diversity in how stable lateral root tip angle is across development (Figure 5 C). While many accessions have a consistent angle of new growth, such as TDr-18 and San2 (Figure 5 C), others change their angle of new growth over time, such as TAA14 and Ting-1 (Figure 5 E).

#### RSA traits are heritable and highly polygenic

In addition to looking at the diversity of RSA traits, we next wanted to use GWA to find potential causal loci. Using the fitted values, we conducted GWA analyses for all nine traits at each time point using the program GEMMA (Zhou and Stephens 2012). In total, 29 SNPs were significant using Bonferroni multiple-test p-value correction at least in one time point (Supplemental Table 3). These alleles also had a minimum allele frequency greater than 0.05. These SNPs highlight regions of the genome with elevated significance and the temporal nature of our data allows us to track the changes in significance over time (Figure 6 A, Supplemental Figure 7). For example, when looking at average angle over time we see an increase in significance, which likely corresponds to the stabilization of the trait (Figure 6 A, Supplemental Figure 7 B). In contrast, we see overall significance decrease for depth-to-width ratios over time (Supplemental Figure 7 B), which likely reflects that growth of root systems becomes constrained at later time points by the physical limits of the rhizotron. We gain confidence in the chromosomal regions identified by tracking the significance over time and observing the slight enrichment of experimental p-values in our Q-Q plot distributions (Supplemental Figure 7).

Out of the 29 identified significant SNPs, four synonymous SNPs can be found within the coding sequence of two genes, eight were in intergenic regions, whereas the majority of SNPs were found in upstream- and downstream regions of genes, potentially affecting expression patterns (Supplemental Table 3). The identification of many significant SNP associations suggests that RSA development is a polygenic trait.

Looking at the two traits with the highest heritability (Supplemental Table 4), average angle (H^2^ = 0.45) and average angle per day (H^2^ = 0.31), we found 12 and 8 significant SNPs, respectively (Supplemental Table 3). Interestingly, both traits revealed one common SNP upstream of a hypothetical protein with 30% protein identity to a phospholipase D alpha 3 (At5g25370) (Supplemental Table 3). Furthermore, the only intragenic SNP for average angle per day was found in an intron of another hypothetical protein with 35% identity to a non-specific phospholipase C2. Phospholipases are lipid-hydrolyzing enzymes that are known to play a role in signaling during plant development, stress responses, and responses to environmental cues (Takáč, Novák, and Šamaj 2019). Our identification of significant SNPs associated with hypothetical proteins similar to phospholipases might indicate a role for these proteins in adjusting root angles to local environments.

Root depth was associated with a SNP upstream of a Cyclin A2;3 (AT1G15570) (Supplemental Table 3). This gene was previously found to play a role in auxin-dependent mitotic-to-endocycle transition that is involved in the transition from cell proliferation to cell differentiation in the Arabidopsis root meristem (Ishida et al. 2010). Thus, alterations in gene expression could influence root depth in certain Arabidopsis accessions.

The GWAS on the depth-width ratio yielded an intronic SNP in a P-loop nucleoside triphosphate hydrolases superfamily protein with Calponin Homology domain-containing protein (Supplemental Table 3). This gene is expressed in the lateral root cap (http://bar.utoronto.ca/eplant/; (Waese et al. 2017)) and changes in expression may influence root gravitropism leading to a more shallow root system. Future work will establish the molecular basis for the SNP-phenotype relationships identified here.

#### RSA traits of the GLO-Roots diversity panel significantly correlate with climatic variables

To gain insight into the relevance of the different root architectures of the GLO-Roots diversity panel in the natural environment, we correlated our nine root traits with published bioclimatic variables (Supplemental Figure 8) (Fick and Hijmans 2017). We see the strongest correlations between these traits and climatic variables related to temperature (Supplemental Figure 8). Specifically, average angle per day shows a notable correlation to mean diurnal temperature range (Pearson r = 0.373, p = 2.56 x 10^-4^) (Figure 6 B). This result indicates that accessions with shallow root system architectures tend to grow in climates that exhibit larger changes in the daily minimum and maximum temperatures. Likewise, convexhull shows a positive correlation to mean diurnal range, albeit less significant (Supplemental Figure 8). These data suggest that plants with shallow roots may actually grow better than those with deep roots in a more variable environment. Clustering of accessions by change in average angle per day over time divides the accessions into three groups: consistently shallow, consistently deep, and intermediate or changing root angle (Figure 6 C, D). Interestingly the consistently shallow and consistently deep accessions segregate from each other in a global map, which is consistent with our finding that climate is likely to be an important variable determining the distribution of accessions with these traits (Figure 6 C).

## Discussion

Implementation of automation expands the number of individuals that can be phenotyped and the phenotypes that can be captured (Gehan et al. 2017). By automating the novel-GLO-Roots system, we created GLO-Bot, a robotic platform that enables unprecedented insight into Arabidopsis root growth over time in soil and at developmental stages rarely observed. Careful design of the system to facilitate automated handling allowed us to maintain physiological relevance while increasing throughput and allowing for novel trait measurements. Along with the automated imaging system, we developed an improved image analysis pipeline for quantification of root system architecture (RSA). The previous image analysis program was semi-automated, and often needed intervention to define regions of interest. While this is feasible for end-point imaging with a small number of samples, it was not scalable with GLO-Bot. For automated image analysis, however, one of the biggest challenges was the combination of maximizing root luminescence signal intensity while decreasing background noise so that GLO-RIAv2 would only analyze true root signal. We saw a strong correlation between our traits extracted via our new image processing pipeline and those that were measured manually, demonstrating that our updates allowed for accurate examination of growth dynamics in both new growth and at a whole root system level. Recent progress in machine-learning is demonstrating the opportunities within the field for segmenting out root and noise and could further improve accuracy (Wang et al. 2019; Smith et al. 2020). Additionally, using transgenic lines homozygous for the luciferase transgene would lead to stronger signal and therefore better root detection. New advances in recapitulating a full bioluminescence pathway *in planta* (Khakhar et al. 2020) could provide an exciting alternative for increasing root luminescence signal and therefore simplifying image segmentation.

The automation of GLO-Roots allowed for imaging larger sample sizes and thus enabled the visualization of root growth of different Arabidopsis accessions over time. A set of six accessions was used to test the robustness of the system and to optimize imaging and watering times in order to extract root traits that reflect true differences in RSA. Nine root traits were automatically extracted through GLO-RIAv2 and showed striking differences even in this small subset of accessions. The GLO-Roots diversity panel enabled the identification of loci associated with variation in the nine root traits measured. However, given the nature of these broad traits, each is likely highly polygenic and, therefore, our ability to detect individual causal SNPs or peaks may be relatively low. Additionally, our relatively small population size (93 accessions) challenges our ability to discover SNPs significantly associated with the traits (Gibson 2012). Nonetheless, we were able to identify significant SNPs for six of the nine root traits, most of which are located up- and downstream of candidate genes and may potentially affect gene regulation. Although a surprisingly rapid linkage decay in *Arabidopsis thaliana* (Nordborg et al. 2005) allows within-gene trait mapping (Atwell et al. 2010; Exposito-Alonso et al. 2018), it will be ideal to follow up the SNPs identified above and other close variants for further analyses, such as accession-specific gene expression and complementation, knock out and reporter gene assays (Ogura and Busch 2015).

The correlation of our nine extracted root traits to published bioclimatic data reveals a significant correlation between average angle to mean diurnal temperature range. Thus, accessions with shallow root system architectures tend to grow in climates that exhibit larger changes in the daily minimum and maximum temperatures reminiscent of desert-like climates. This fits with the known pattern that many plant species growing in the desert show rather shallow root systems with advantages including reduced energy input, increased ability to capture moisture from precipitation, and high nutrient availability in upper soil layers, which may improve survival in a more varied climate (Pierret et al. 2016; Schenk 2008; Ogura et al. 2019).

Surprisingly, we found low or some non-significant correlations between the nine root traits and total annual precipitation, a factor we had initially used to select the natural populations used in this study. This finding could be due to a variety of factors underlying the design of our study including the limited number of accessions analyzed, the specific selective pressures acting on Swedish Arabidopsis populations, where the majority of our characterized accessions originate (Long et al. 2013), and the well-watered conditions that we used to grow our plants. It may also be that annual precipitation is less important as a selective pressure acting on Arabidopsis root architecture compared to temperature. Indeed vapor pressure deficit (VPD), which is the driving force determining the rate of water loss during transpiration (Fricke 2017), is exponentially related to temperature and modern crop plants appear to be particularly susceptible to elevated VPD (Lobell et al. 2014). Since we assayed root architecture under well watered conditions, the architectural features we uncovered may be primarily relevant to seasonal conditions where water is sufficient, but where temperature fluctuations create a physiological demand for water that requires an expansive, but shallow root system. Thus, surveying our accessions in conditions similar to their native environment or in conditions replicating environmental stresses will help expand our understanding of architectural traits related to climate.

The climate models recently published by the Intergovernmental Panel on Climate Change (Masson-Delmotte, V., P. Zhai, A. Pirani, S. L. Connors, C. Péan, S. Berger, N. Caud, Y. Chen, L. Goldfarb, M. I. Gomis, M. Huang, K. Leitzell, E. Lonnoy, J.B.R. Matthews, T. K. Maycock, T. Waterfield, O. Yelekçi, R. Yu and B. Zhou (eds.) n.d.) predict severe trajectories with extreme weather events and thus higher climate variability at most geographic locations. Our findings suggest that environmental variability – particularly variability in temperature – rather than environmental intensity, creates a physiological condition where root angle traits matter. By taking a physiologically-relevant approach to observe these responses in a model system, we hope to contribute to predicting how the invisible part of plants will react to an ever warming climate, both in agricultural and in natural ecosystems.

## Data availability

GLORIAv2 is available through Zenodo, DOI: https://doi.org/10.5281/zenodo.5574925

Image analysis pipelines and scripts are available through Zenodo, DOI: https://doi.org/10.5281/zenodo.5708430

RShiny App for exploring root system architecture of accessions is available through Zenodo, DOI: https://doi.org/10.5281/zenodo.5708422

Imaging data and images are available through Zenodo, DOI: https://doi.org/10.5281/zenodo.5709009

Previously published datasets used: WORLCLIM2: Fick SE, Hijmans RJ, 2017, https://worldclim.org/, https://doi.org/10.1002/joc.5086

## Acknowledgements

Work in the JRD lab was funded by the U.S. Department of Energy’s Office of Biological and Environmental Research (DE-SC0008769 and DE-SC0018277) and the Carnegie Institution for Science Endowment. TSL was funded by the National Science Foundation Graduate Research Fellowship and the National Institutes of Health Predoctoral Training Grant (T32GM007276). HL was funded by a German Research Foundation (DFG) research fellowship (LI 2776/1-1). MEA is funded by the Carnegie Institution for Science, an NIH Early Investigator Award (1DP5OD029506-01), and a DOE BER grant (DE-SC0021286). GL is funded by the German Research Foundation under Germany’s Excellence Strategy, EXC-2070 - 390732324 (PhenoRob). Part of the computational analyses were done at the Carnegie High-Performance Computing clusters *Memex* and *Calc*.

We thank Peter Sand and Zoltan DeWitt from Modular Science for the design and maintenance of GLO-Bot. We thank Wolfgang Busch for advice on the initial selection of Arabidopsis accessions. We thank Chris Ballard from Sutter Instrument for maintenance of GLO1. We thank Theo van de Sande and Ismael Villa for maintenance of the growth room. We thank Michael Raissig and Josep Vilarrasa-Blasi for constructive suggestions on the manuscript.

## Competing interests

The authors declare that no competing interests exist.

## Supplemental material

**Supplemental Figure 1:**
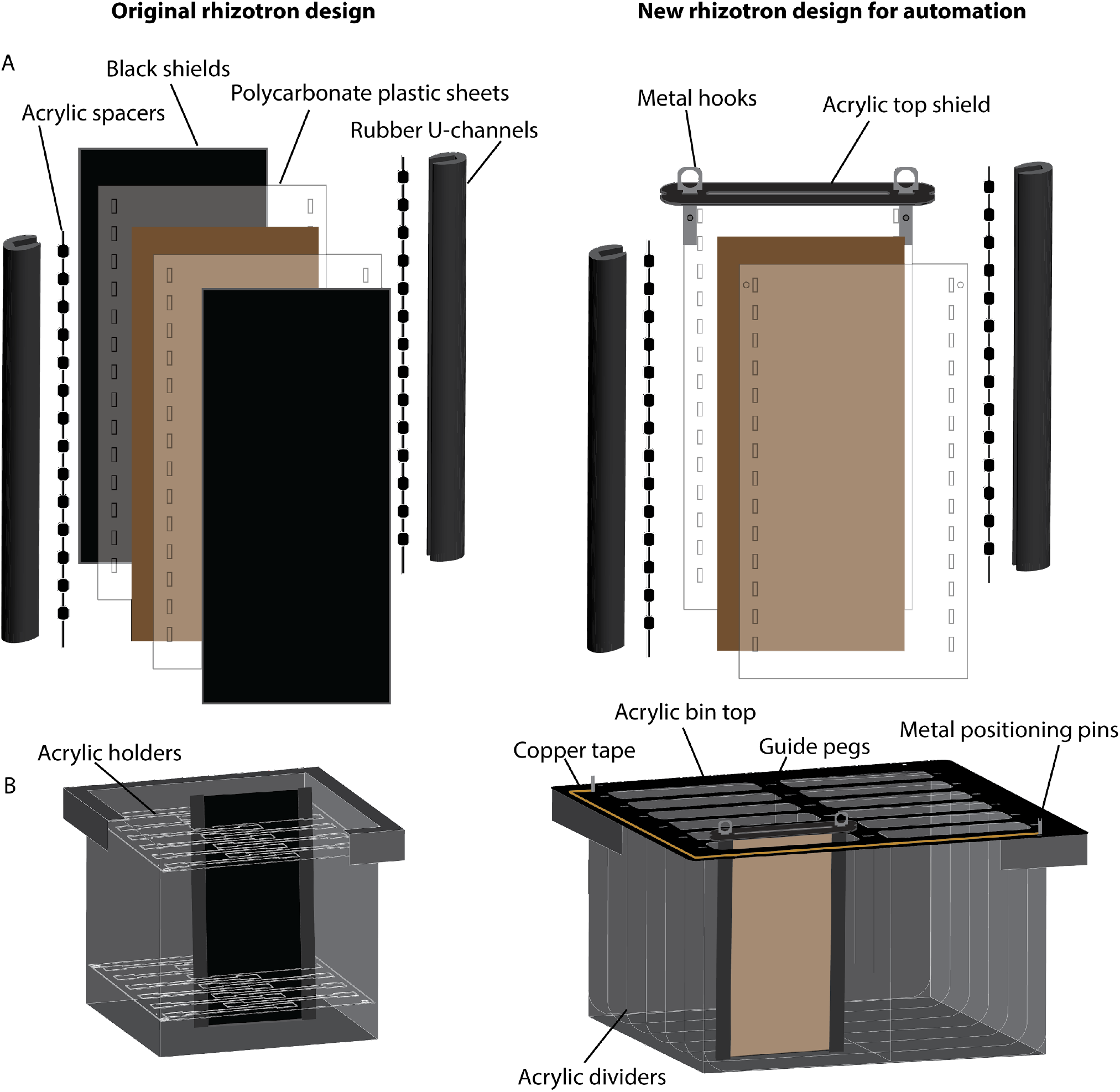
Detailed schematic of rhizotron design for automation. (A) Rhizotron design changes between the original rhizotron design published in Rellán-Álvarez et al., (2015) (left) and new rhizotron for automation (right). The new design uses an acrylic top shield secured in place by metal hooks instead of the black shields to protect roots from light. (B) This light protection is possible since rhizotrons hang in the bin rather than stand in the bin. Guide pegs adhered to the bin top help the robotic arm position pick-up and drop off the rhizotrons while acrylic dividers within the bin provide additional light protection for neighboring rhizotrons if one rhizotron is removed from the bin. Copper tape and metal positioning pins are used by the robotic arm to detect the position of the bin and calculate the exact position of each rhizotron.

**Supplemental Figure 2:**
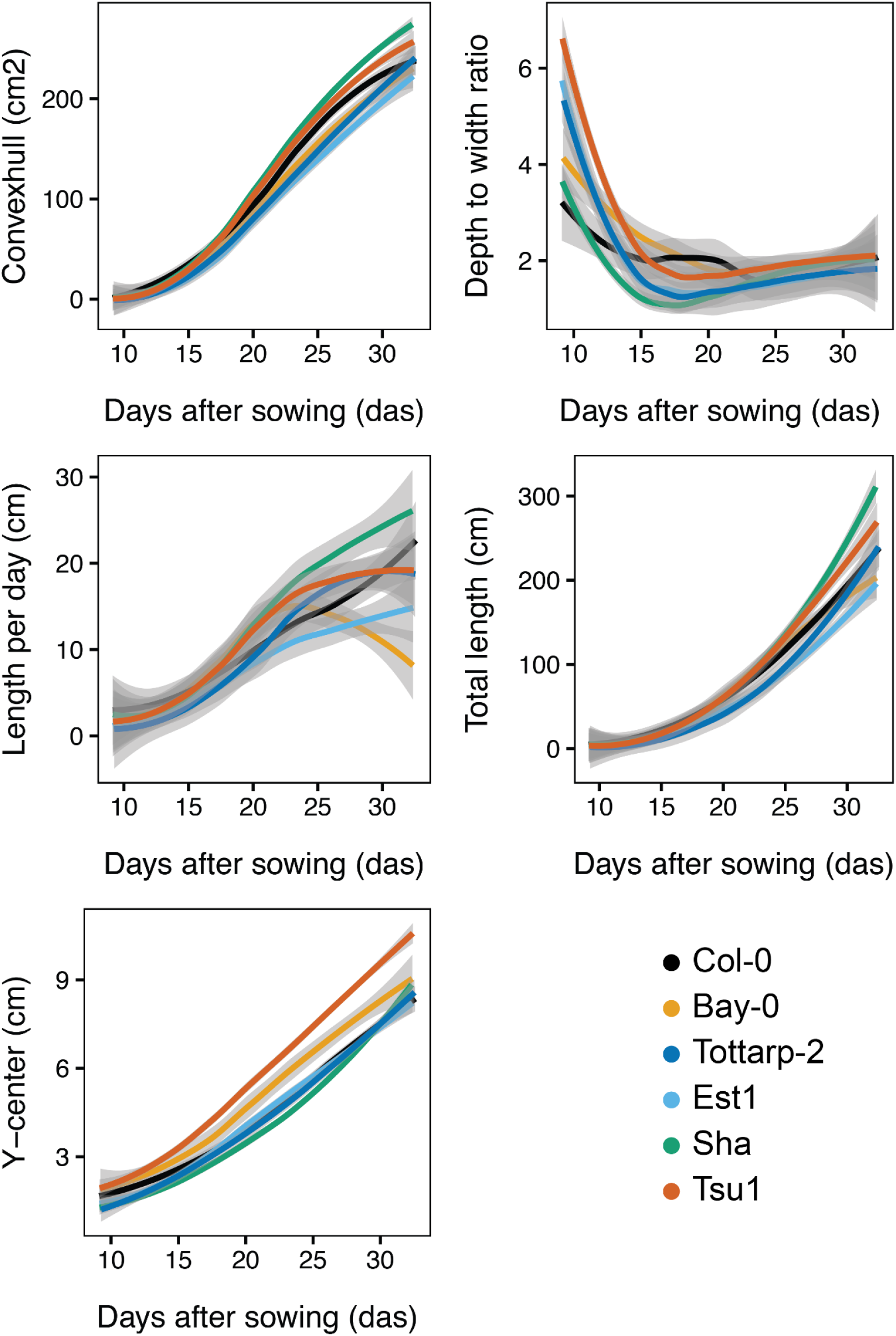
RSA traits over time for the six accessions. Five additional root system traits for the six Arabidopsis accessions growing over time from 9 to 31 DAS. Trends were calculated using Loess smoothing (n = 12-15 plants at each time point). 95% confidence interval shown by grey shading.

**Supplemental Figure 3:**
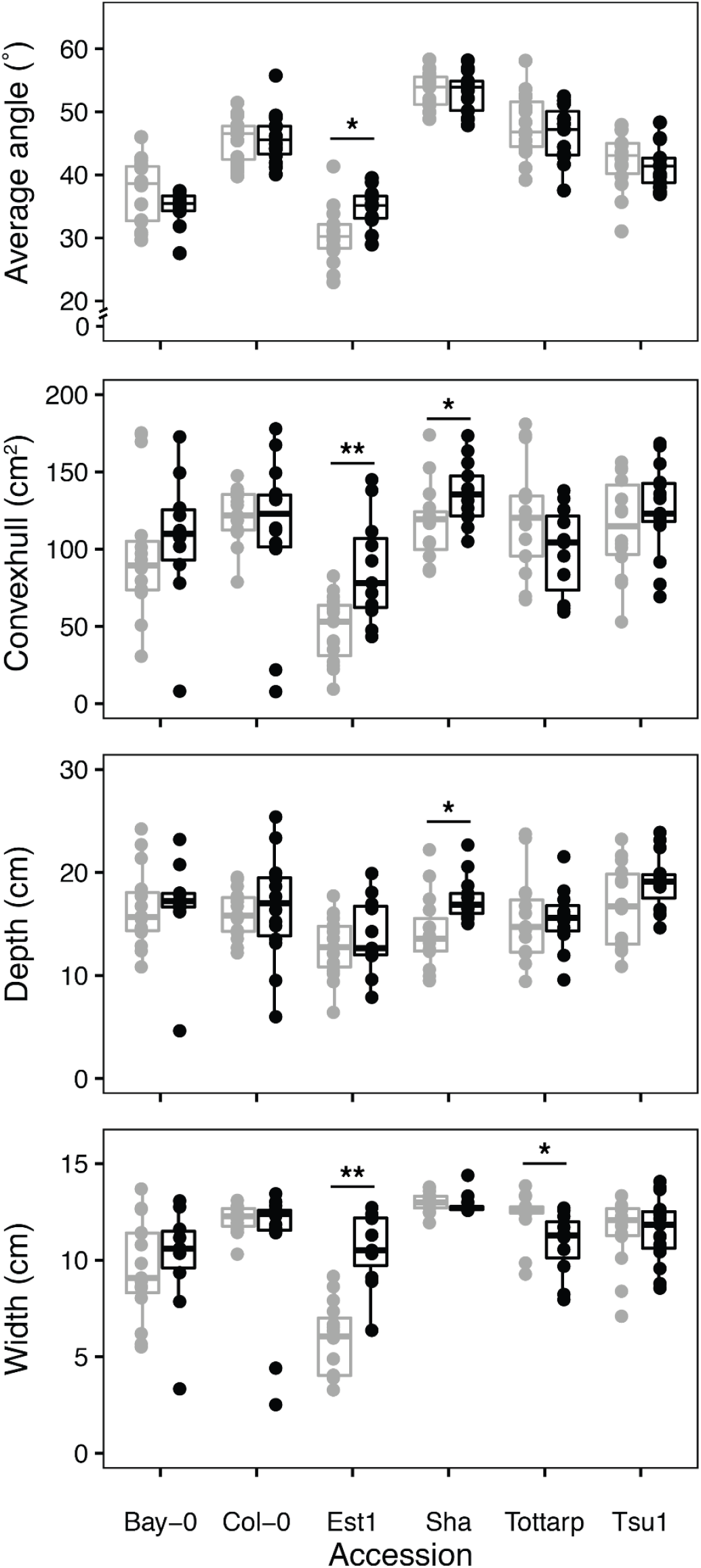
Continuous versus single time-point imaging. Comparison between four traits measured on plants imaged and handled only once at 21 DAS (grey) and plants that were imaged and handled every day up until 21 DAS (black). Comparisons are shown as boxplots (n = 12-15 plants). Box boundaries represent the 25% to 75% percentile range and median value is indicated by the black line. Points beyond whiskers extend beyond 1.5 times the interquartile range. Asterisks indicate significance: * p < 0.05, ** p < 0.01

**Supplemental Figure 4:**
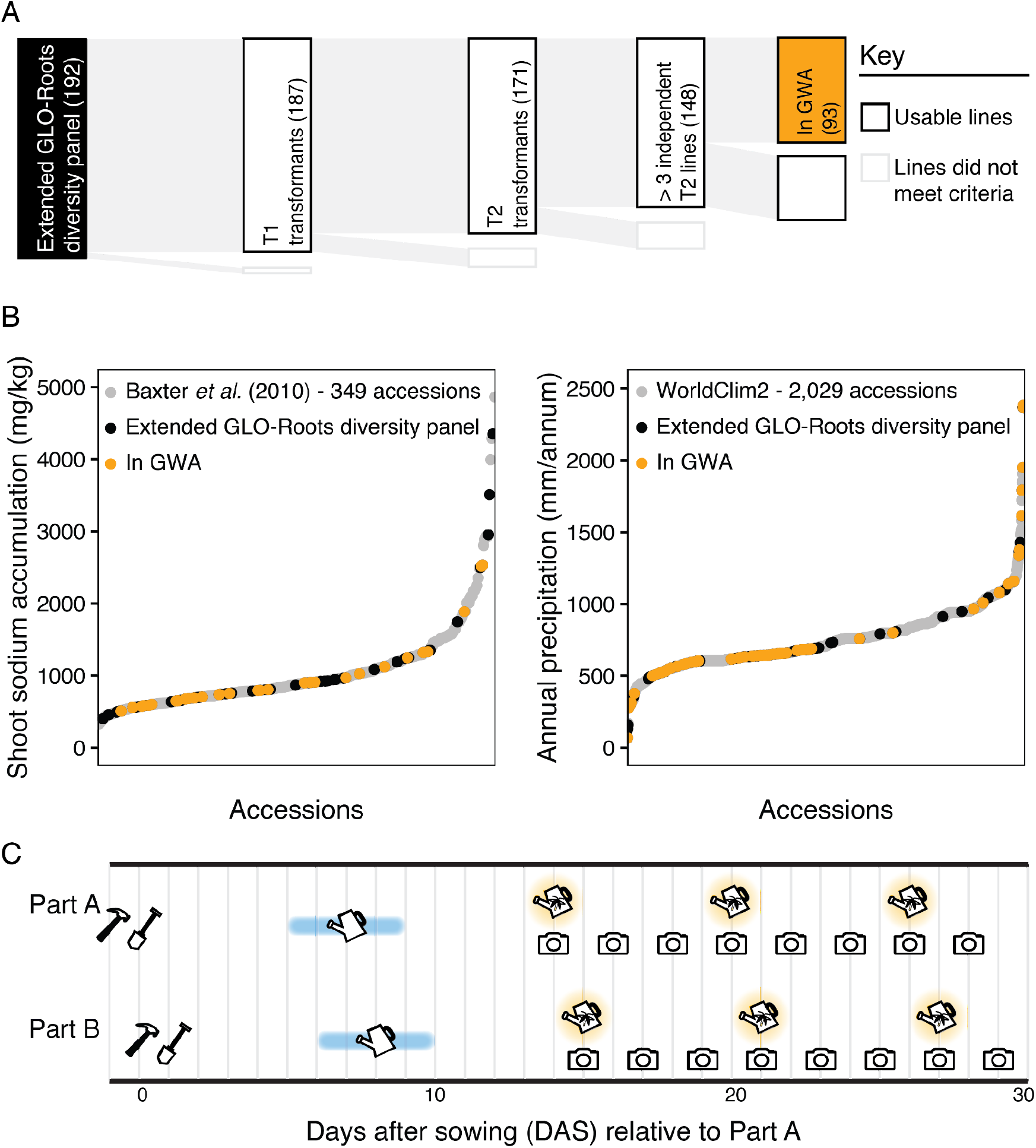
Overview of the GLO-Roots diversity panel. (A) Tracking of the transformation of 192 Arabidopsis accessions and final selection for the genome wide association (GWA) population. (B) Both the extended GLO-Roots diversity panel and the GLO-Roots diversity panel used in the GWAS capture the published distribution of variation found amongst larger populations. (C) Schematic of GLO-Bot growth for the diversity panel including the building (hammer), sowing (shovel), watering (blue watering can), luciferin watering (yellow firefly watering can), and imaging (camera), timeline for one replicate.

**Supplemental Figure 5:**
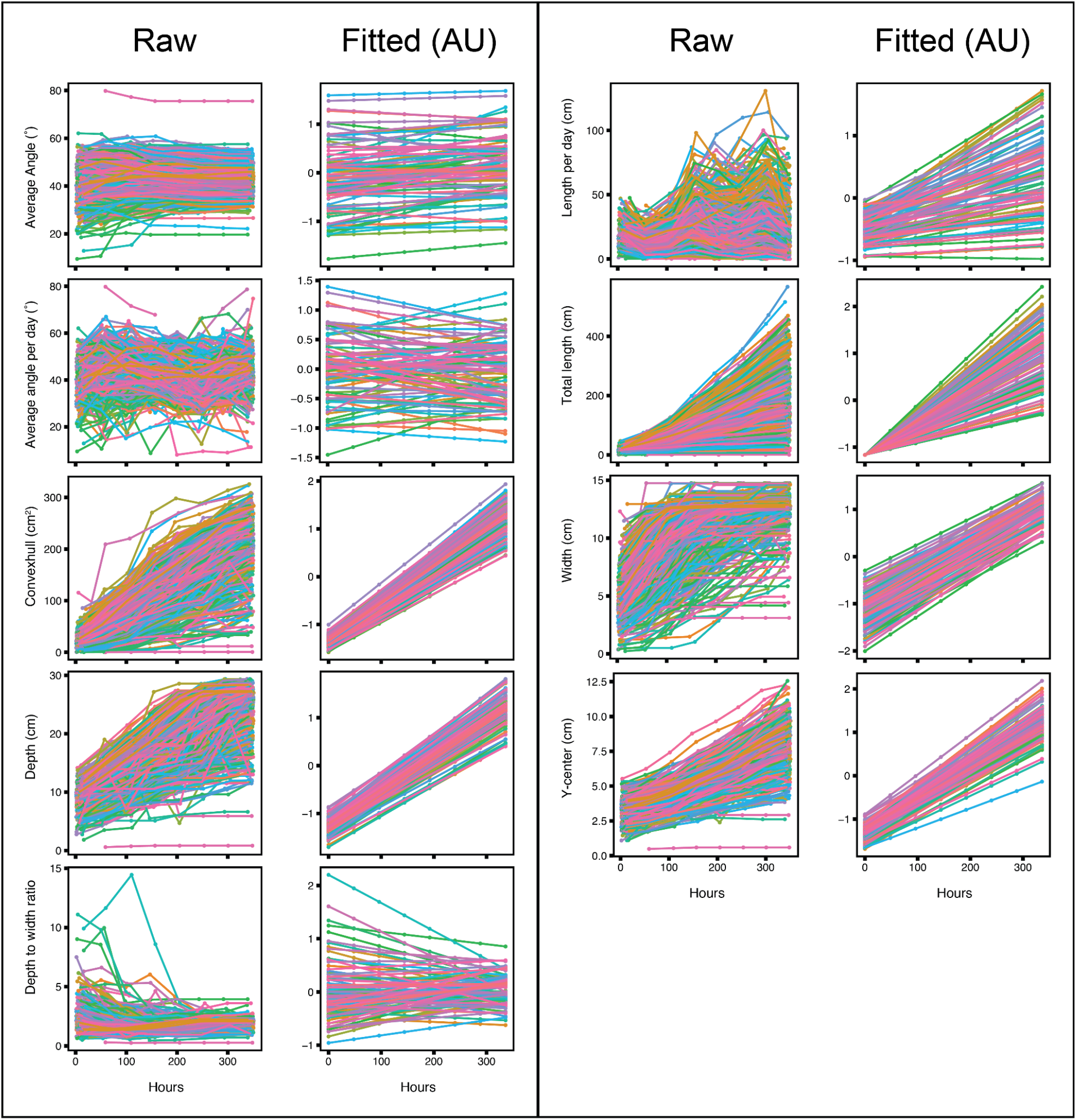
Raw vs. breeding values. Raw (left) and fitted values (right) for all nine root traits. Colored by accession. Time represented in hours after the start of imaging: Hours 0, 48, 144, 192, 240, 288, and 336 are equivalent to 14, 16, 18, 20, 22, 24, and 28 DAS, respectively.

**Supplemental Figure 6:**
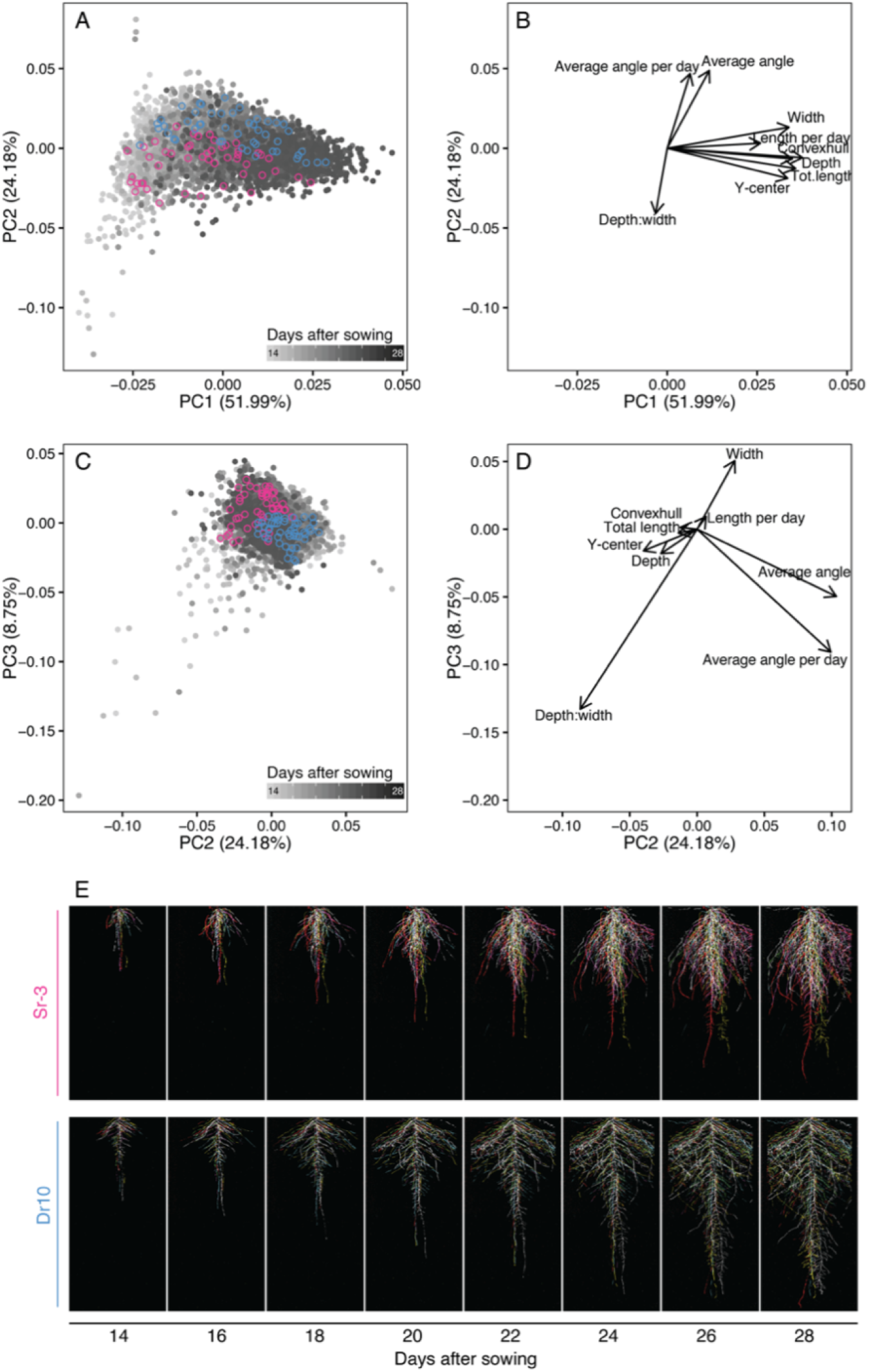
PCA for nine root traits describing RSA. (A-B) Principal component analysis (PCA) of raw RSA traits for the 93 accessions in the GLO-Roots diversity panel shows time and traits correlated with time as the primary axis of variation (PC1). (A-D) Angle and depth to width ratio are negatively-correlated along PC2 and distinguish accessions with steep vs. shallow angles. (A, C) Two accessions, one with steep angles (pink) and the other with shallow angles (blue) are highlighted. (E) Composite images of these two accessions growing over time. Six root systems overlaid on top of each other with root system color indicating replicate.

**Supplemental Figure 7:**
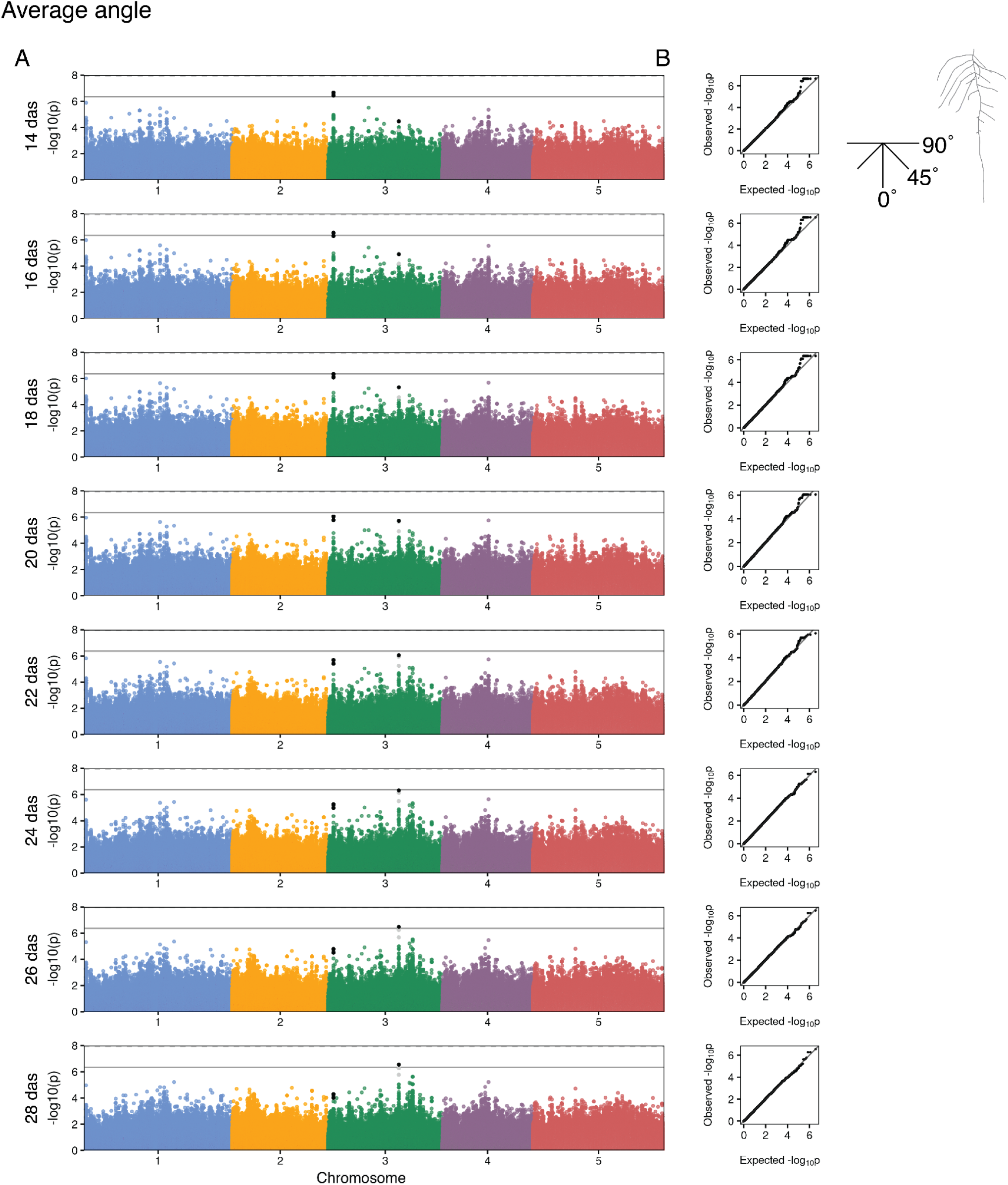
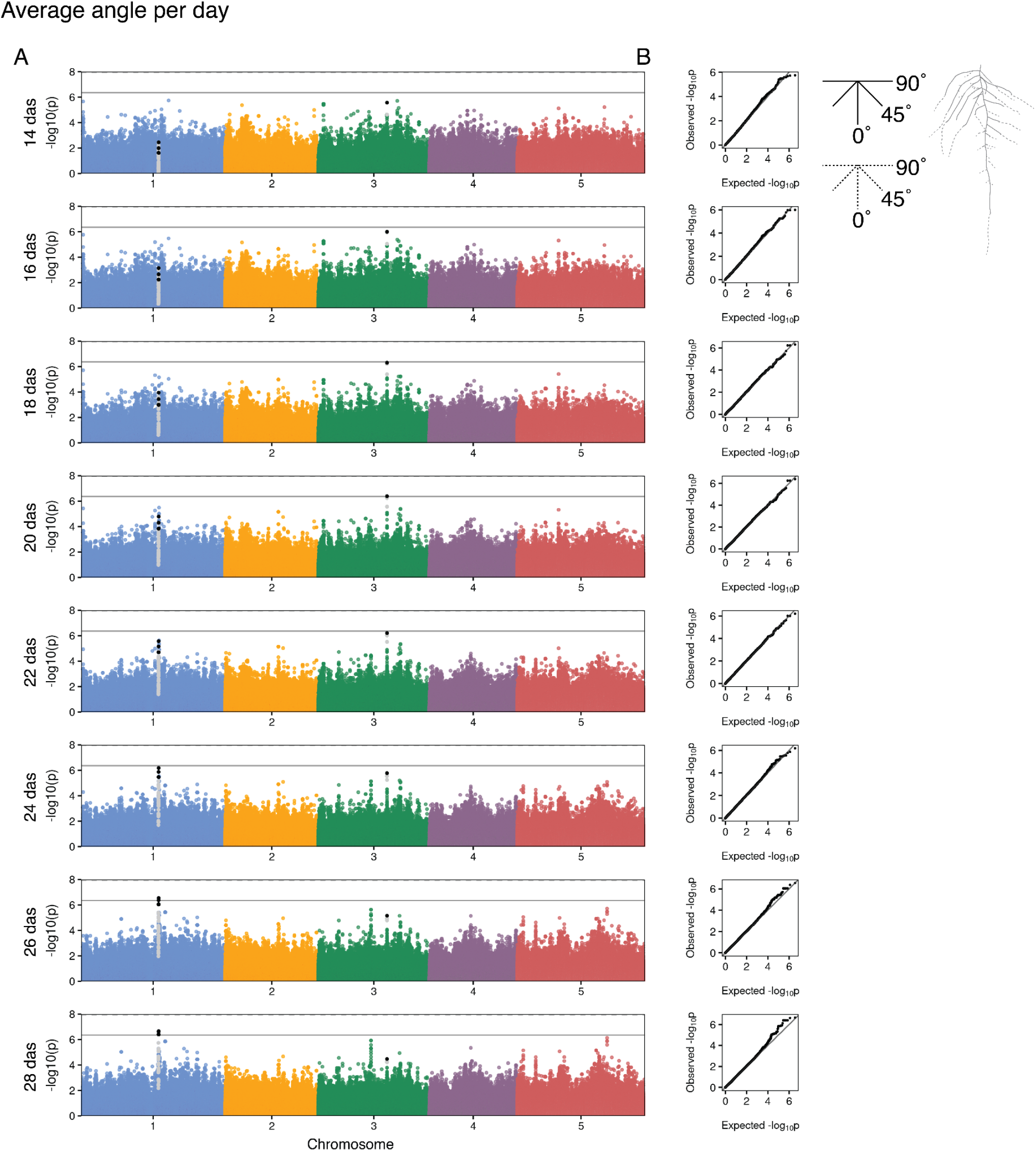
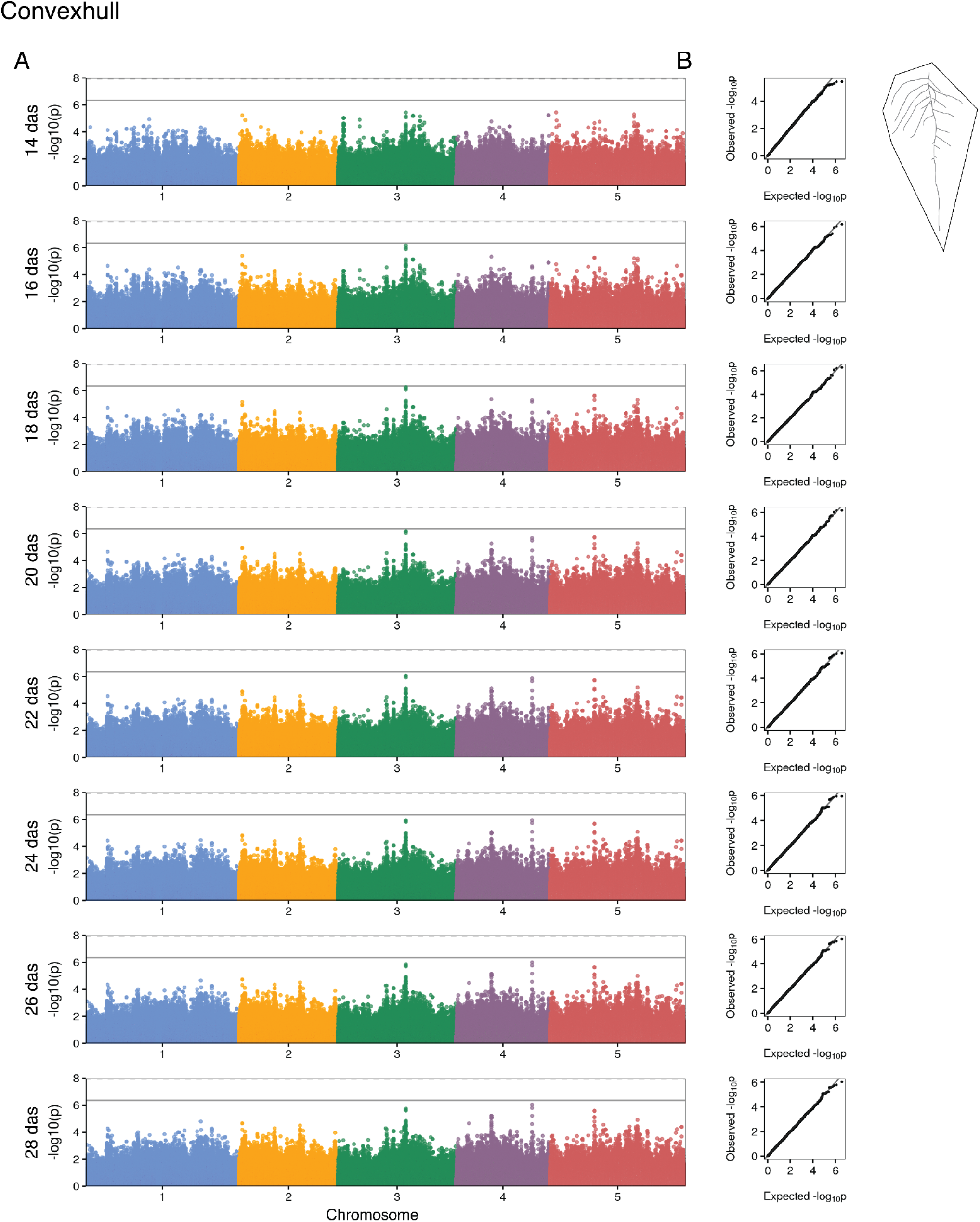
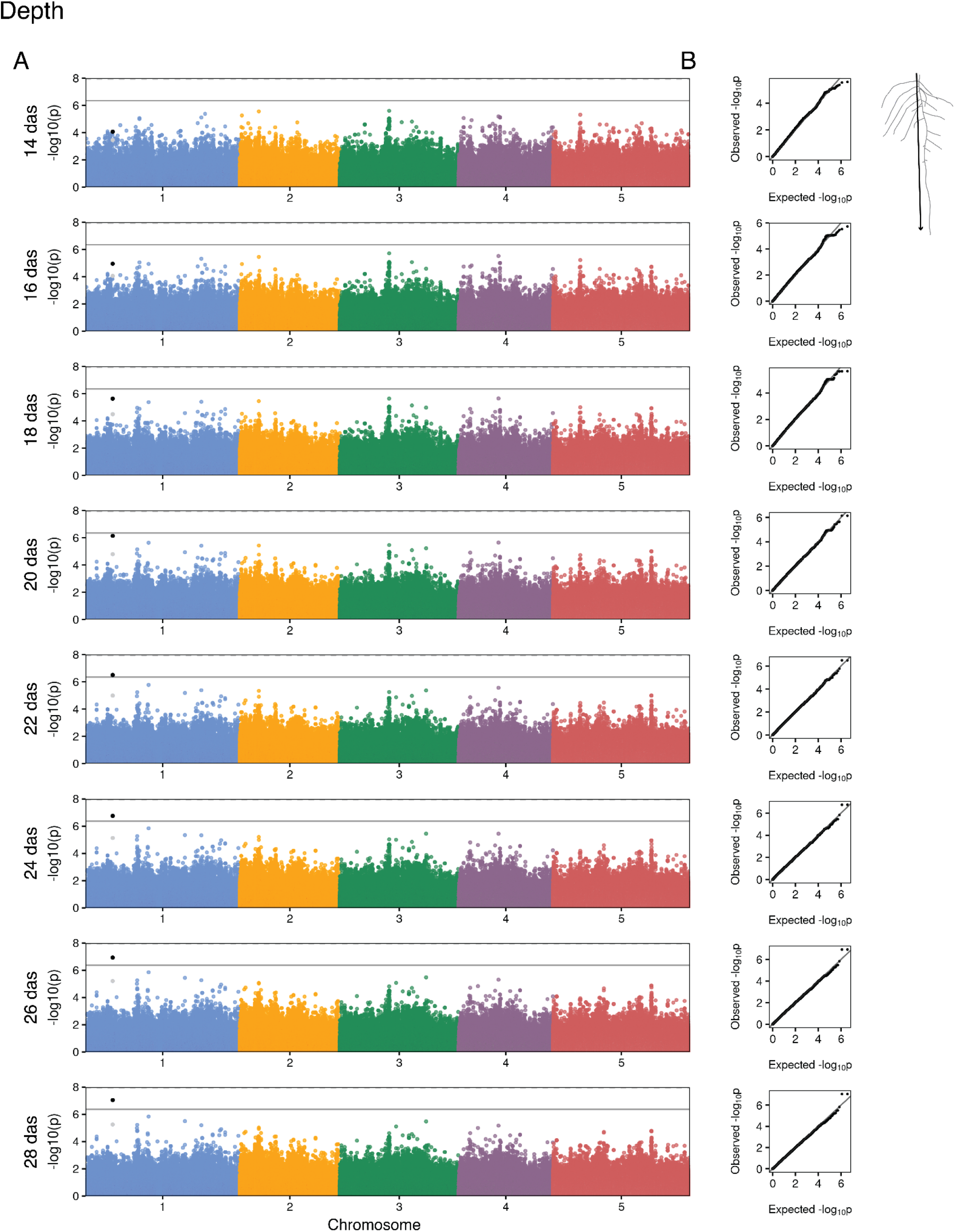
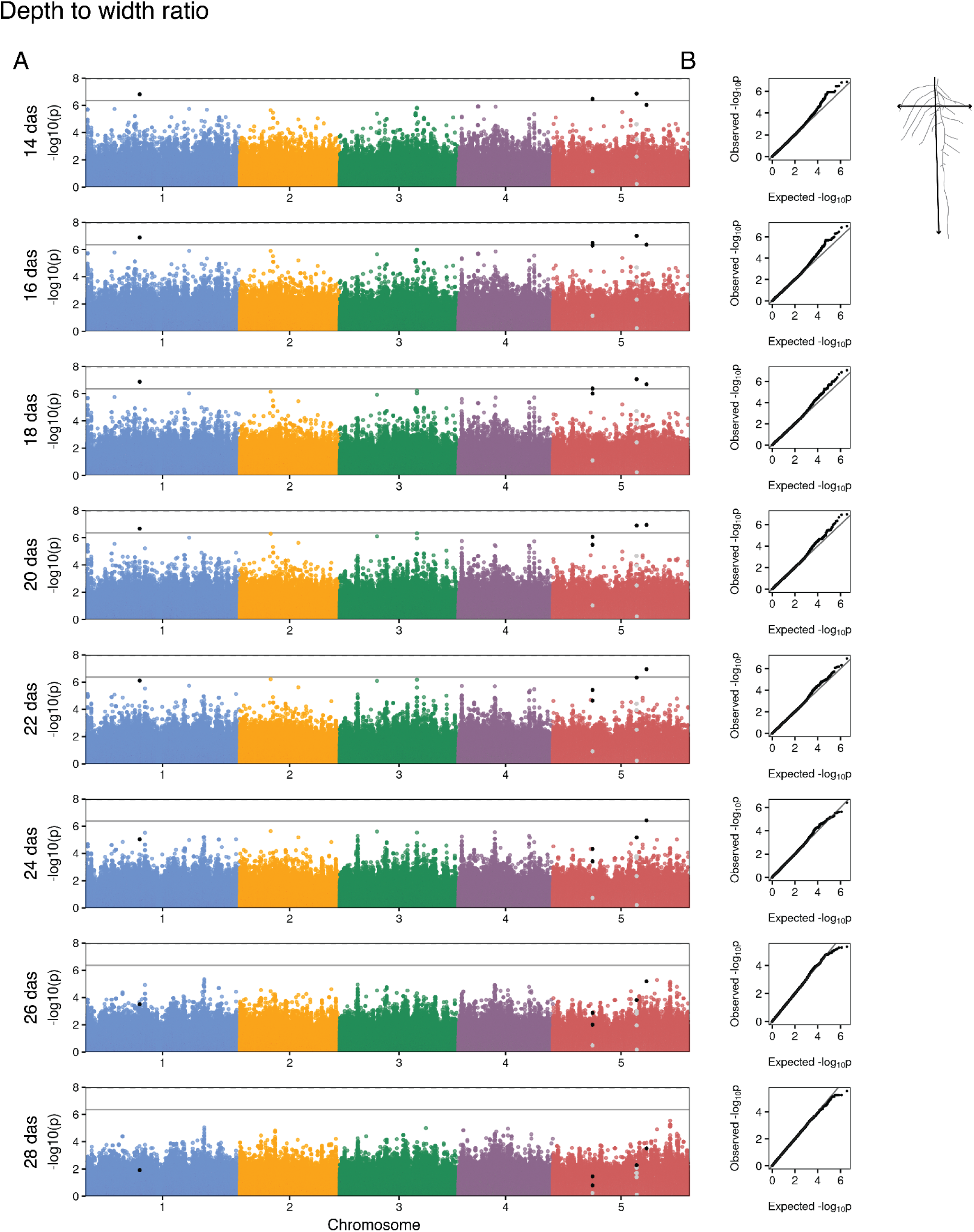
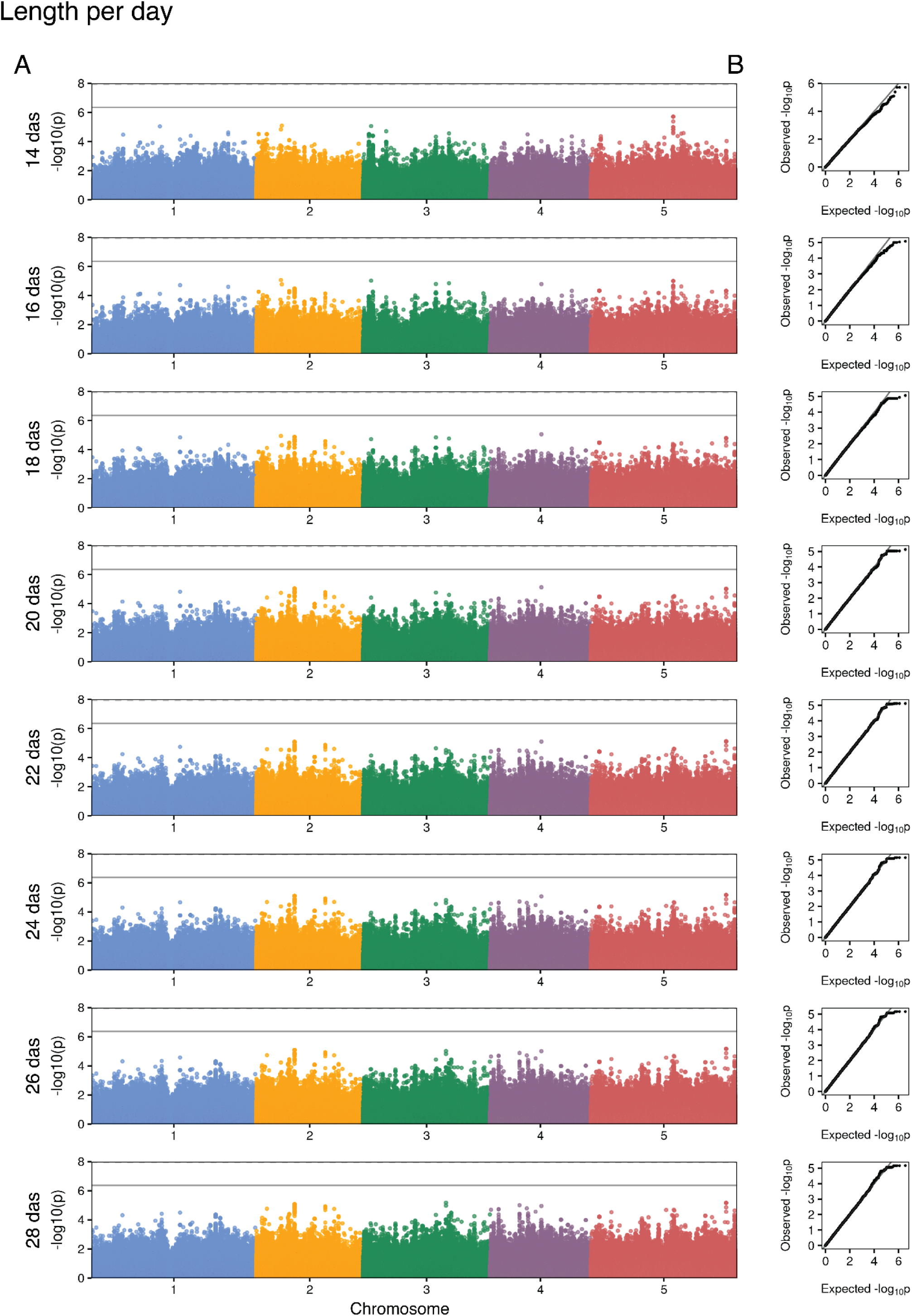
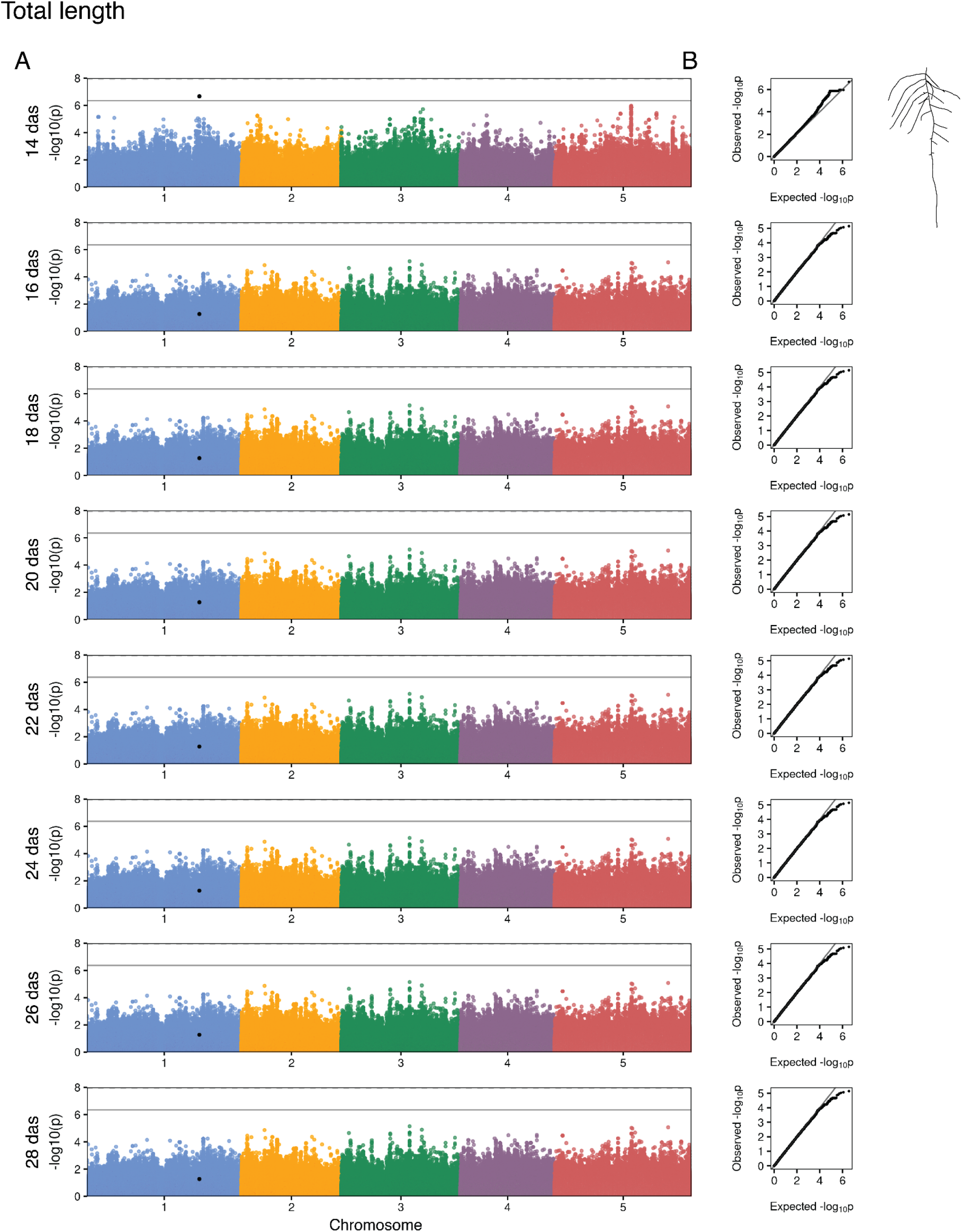
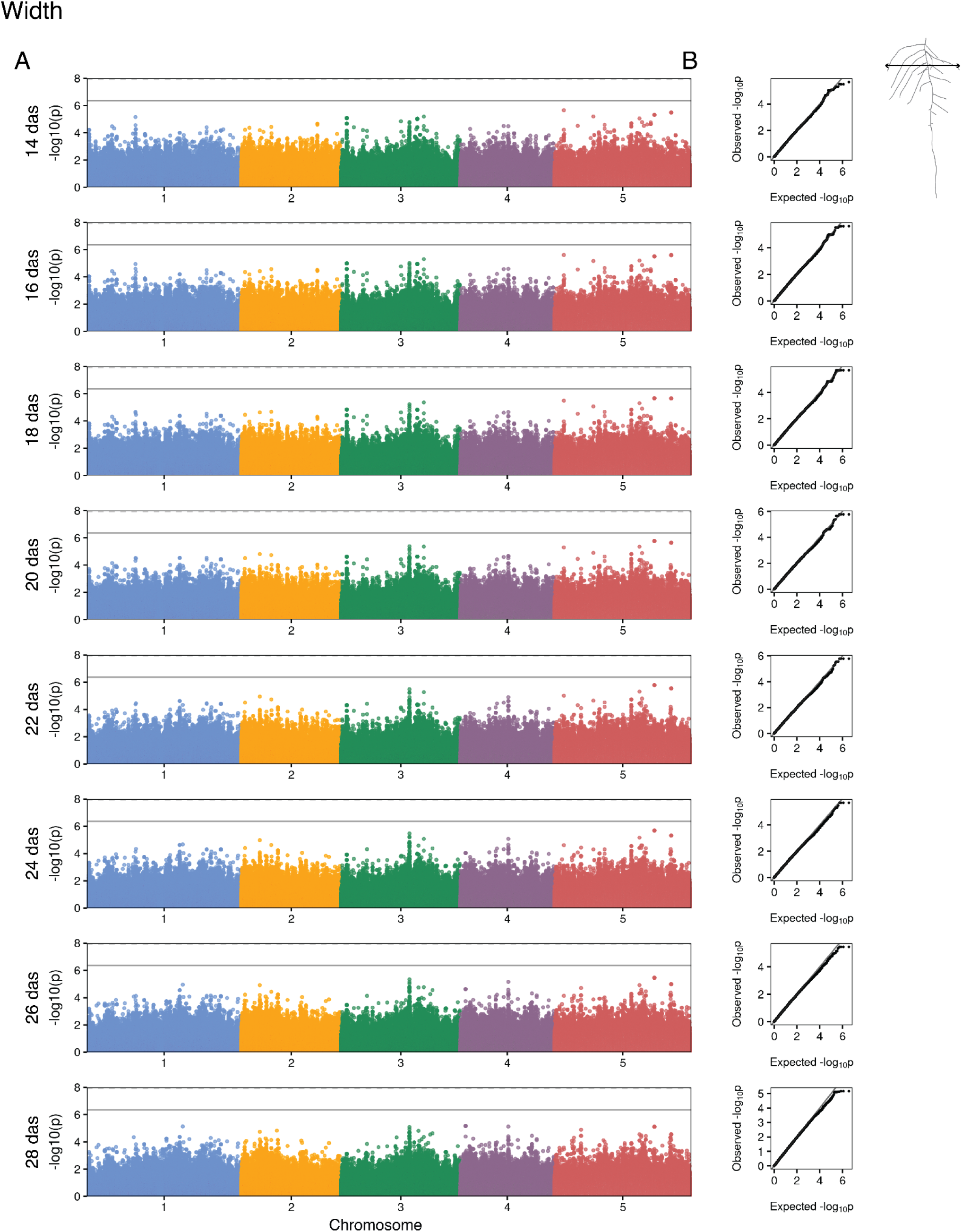
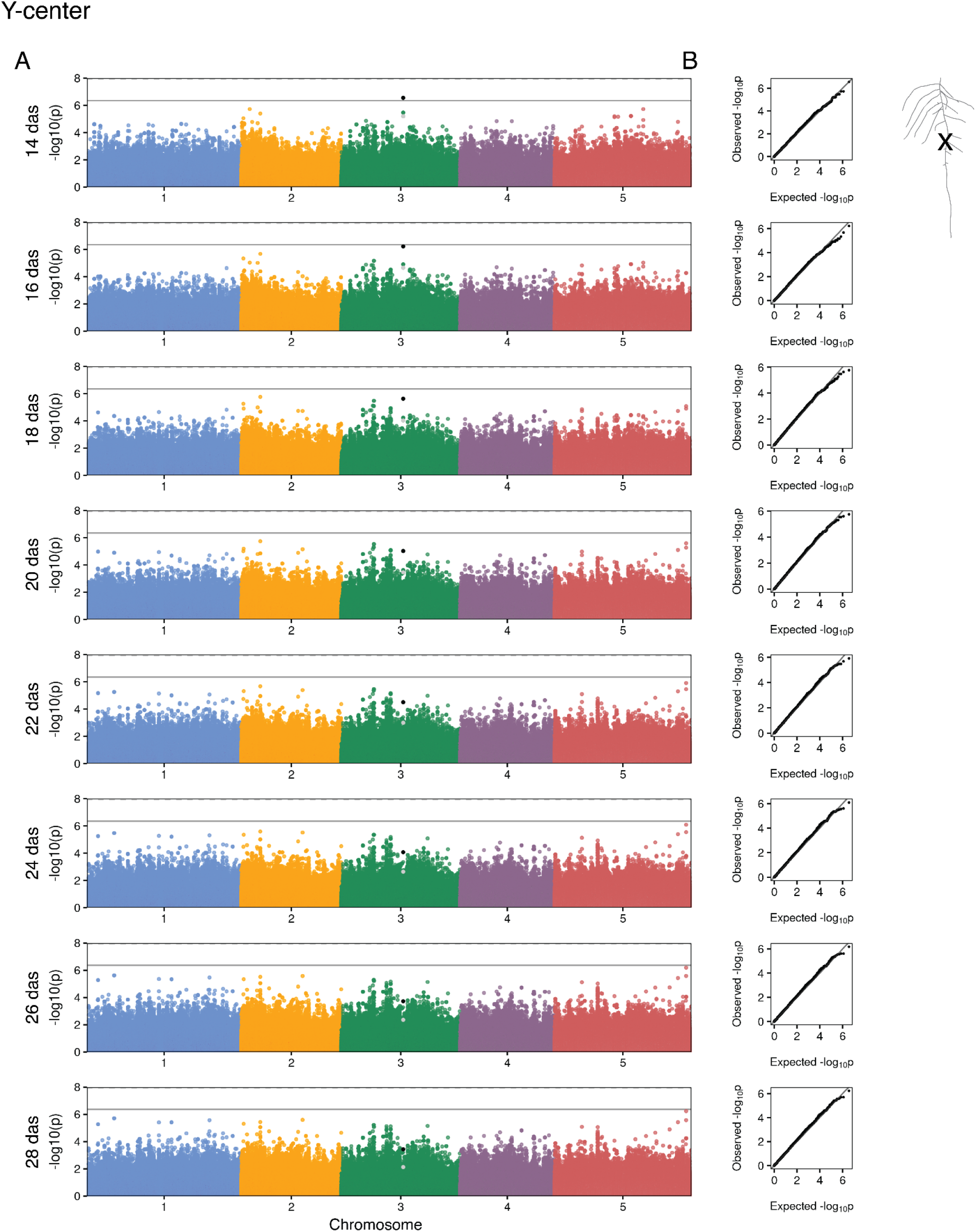
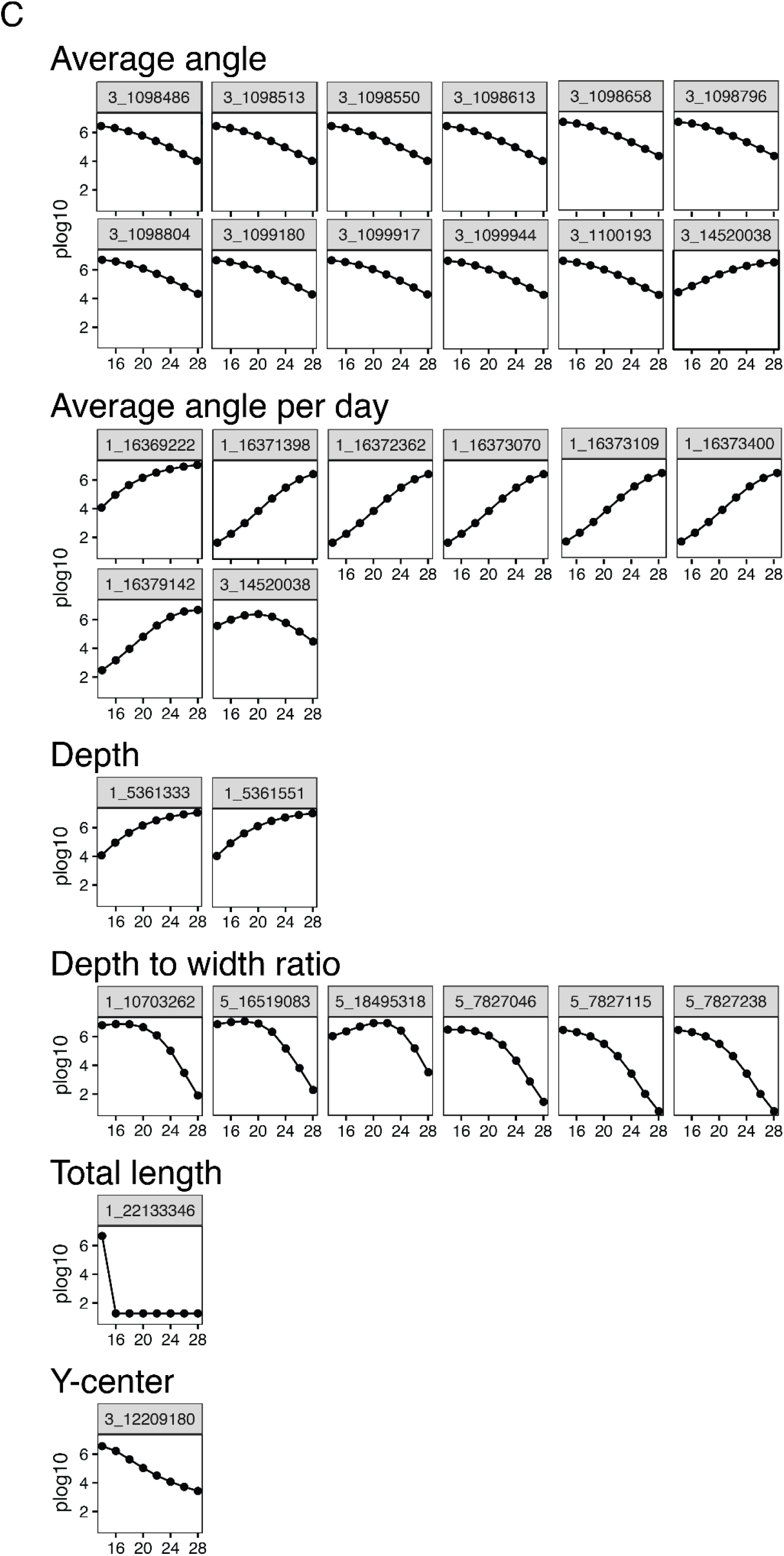
Manhattan plots and SNP development of all nine RSA traits. (A) Manhattan plots of GWA run on each of the 9 RSA traits at each time point with corresponding Q-Q plots (B). SNPs highlighted in black pass the threshold, grey indicates SNPs in linkage disequilibrium (LD). Genes in this region are reported in Supplemental Table 3. (C) SNP development of the 29 unique SNPs that pass the Bonferroni threshold at least once throughout the time series.

**Supplemental Figure 8:**
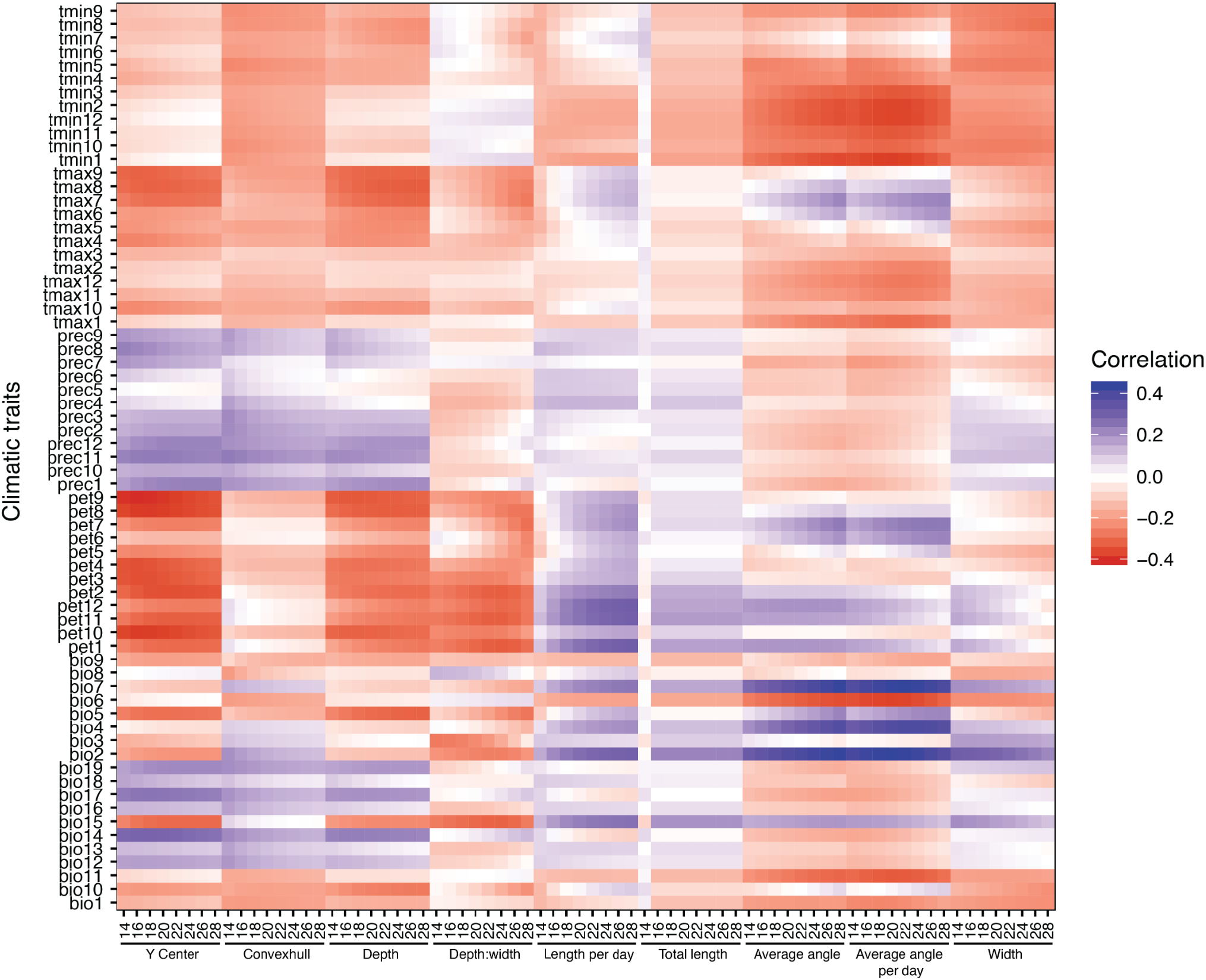
Correlations between average climatic variables and root traits. Pearson correlations between average climatic traits and root traits. We used minimum and maximum temperature for each month (1-12), total precipitation per month, and their composite Bio1-19 were averages in the period 1960-1990 (Fick and Hijmans 2017). Potential EvapoTranspiration (pet) was calculated from the same data as described in (Exposito-Alonso et al. 2019).

Supplemental Video 1: GLO-Bot running

Supplemental Video 2: RSA growth of six Arabidopsis accessions over time

Supplemental Video 3: RSA growth of GLO-Roots diversity panel over time

**Supplemental Table 1:** Analyzed root traits.

| Trait | Units | Description |
| --- | --- | --- |
| Depth | cm | Deepest point identified in root image |
| Width | cm | Left most point detected in root image subtracted from right most point |
| Convexhull | cm<sup>2</sup> | Area within bounding polygon |
| Average angle | Degrees (°) | Length-weighted average angle with respect to gravity of the entire root system such that 0 is straight down and 90 is straight out to the side |
| Average angle per day | Degrees (°) | Length-weighted average angle with respect to gravity of only new growth on a given day |
| Total length | cm | Sum of lengths of all root segments |
| Length per day | cm | Sum of all lengths of new growth root segments on a given day |
| Depth to width ratio | - | Root depth divided by root width |
| Y-center | cm | Vertical centered location of root mass |

**Supplemental Table 2:**
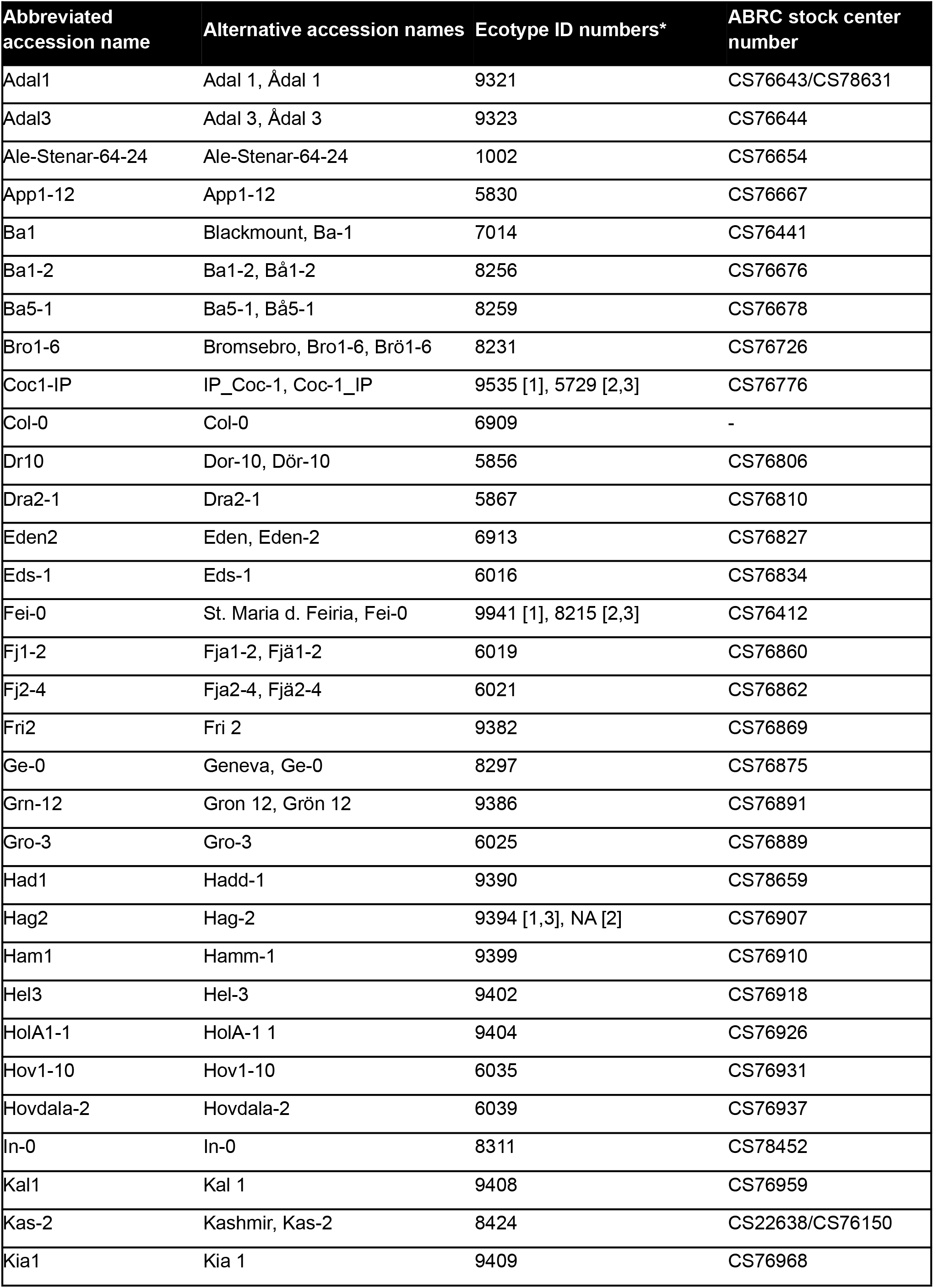
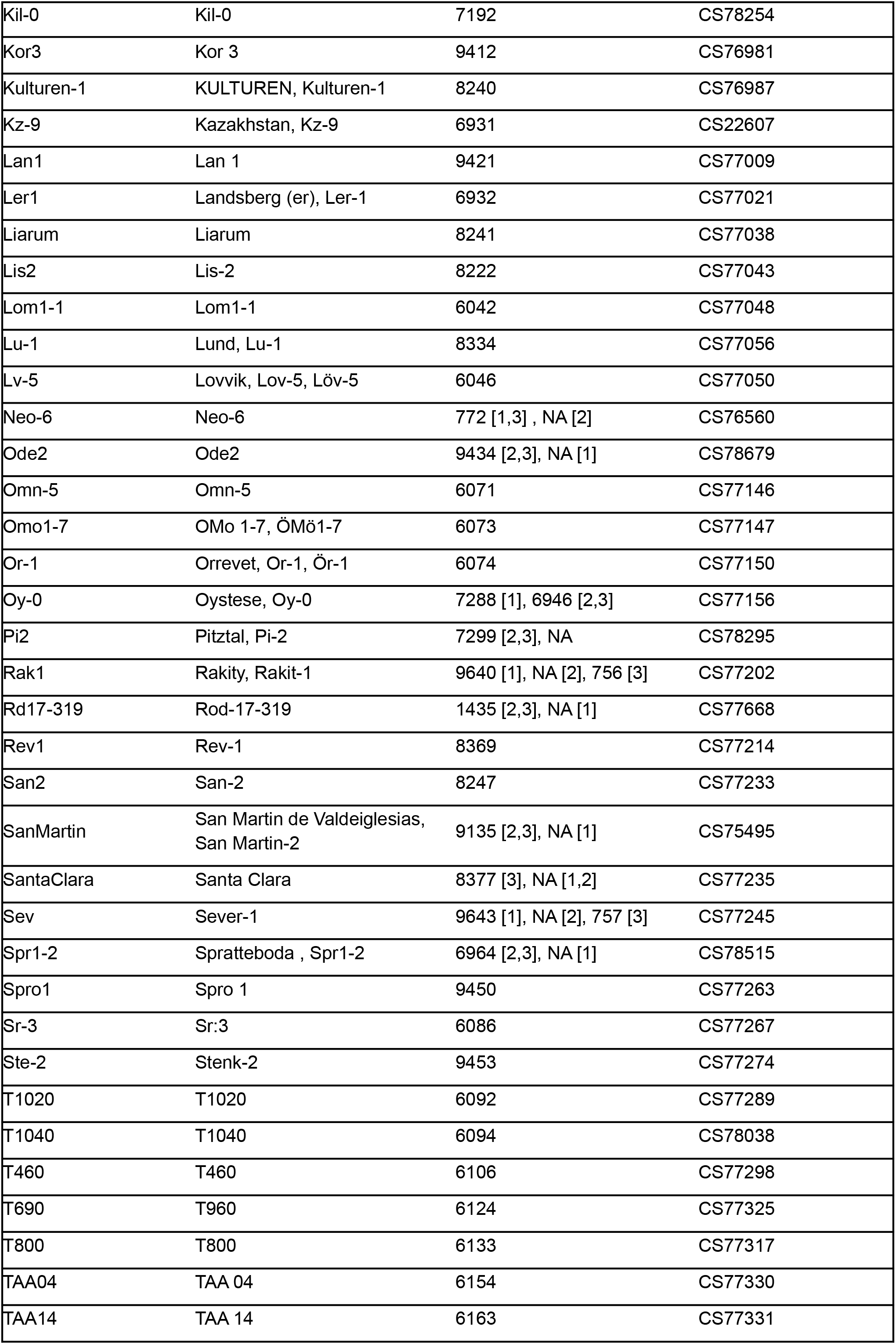
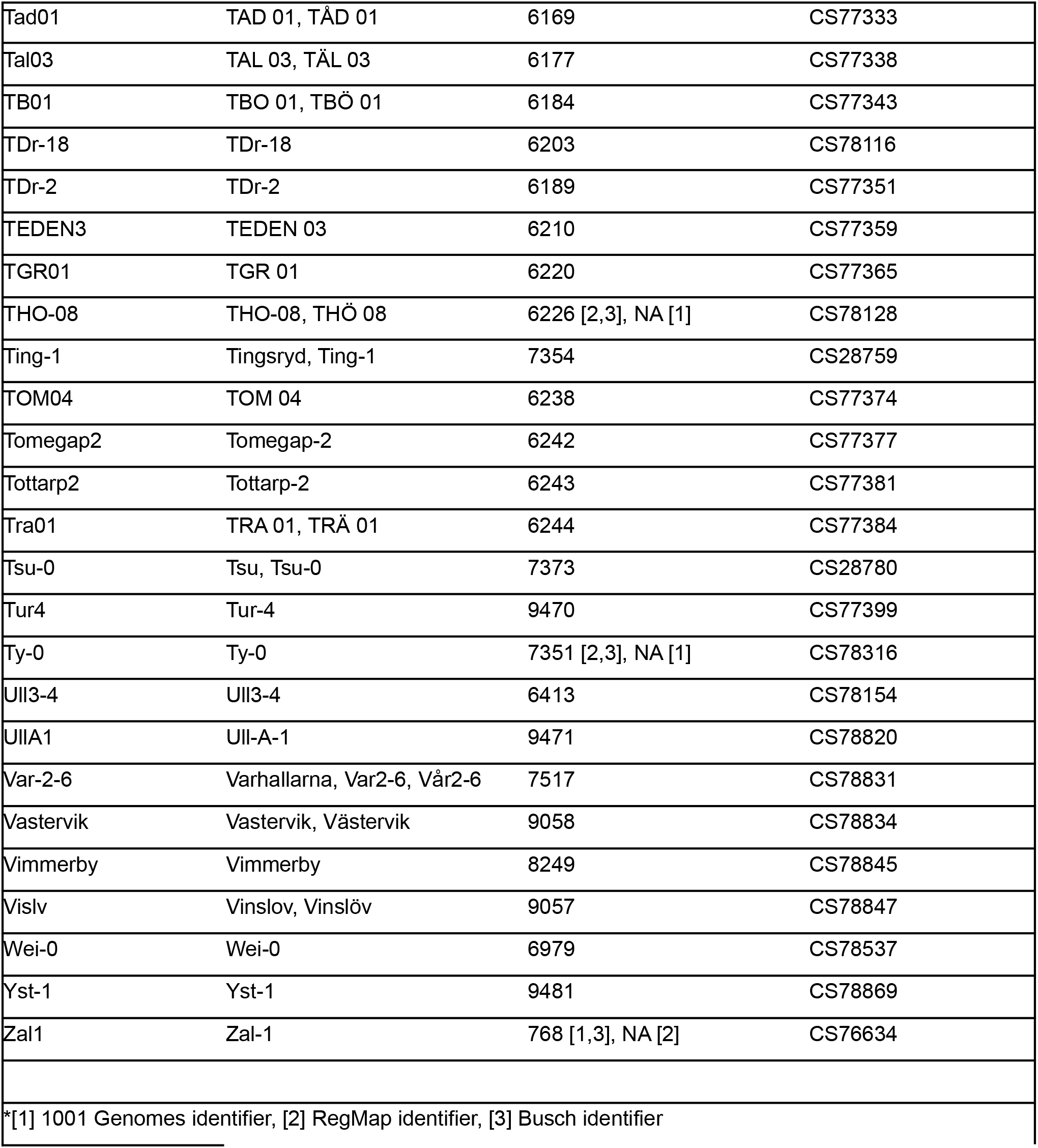
Accessions used for GWAS.

**Supplemental Table 3: SNP list of 29 significant SNPs**

See Excel data table in supplementary materials.

**Supplemental Table 4:** Broad-sense heritability of traits.

| Trait | Heritability |
| --- | --- |
| Average angle | 0.4542113 |
| Average angle per day | 0.3062975 |
| Depth to width | 0.2856909 |
| Width | 0.2787126 |
| Y-center | 0.1165843 |
| Depth | 0.1096065 |
| Length per image | 0.07867771 |
| Convexhull | 0.06618282 |
| Total Length | 1.19E-05 |

## Materials and methods

### Growth system

#### Rhizotron design

As described in Rellán-Álvarez et al., 2015, rhizotrons were built with two sheets of 1/8” clear abrasion resistant polycarbonate plastic (Port Plastics, Portland, OR, Product: Markolon®AR) and were water jet cut (AquaJet LLC, Salem, OR) into 15 cm x 30 cm rectangles with 14 small gaps running down each side (clear_sheets.dxf). Two spacers (spacers.eps) were laser cut (Stanford Product Realization Lab) from 1/8” black cast acrylic (TAP plastics, Mountain View, CA and Calsak Plastics, Hayward, CA). Two rubber U-channels were cut into 29 cm long strips from Neoprene Rubber Trim, 5/16” Wide x 23/32” High Inside (McMaster-Carr, USA, Part Number: 8507K33). Additional adaptations for automation included: beveling the top of each polycarbonate sheet to a thin 45 ° angle using a belt sander. Half of the polycarbonate sheets had 2.5 mm holes drilled in 5/8” down from the top edge and 3/16” in from each side. The small holes were used to screw a 1/16” aluminum water jet cut hook, outer dimensions 18 mm x 57 mm with a 13 mm diameter circle cut out (rhizotron-machine-hooks-13mm.pdf), to both sides of the sheet using 2 x 4 M3 stainless steel socket cap screws (McMaster-Carr, USA, Part Number: 91292A109). Not only did the hooks provide a way to pick up the rhizotron, they also were threaded through a rhizotron top (rhizo-bin-top-enlarged.pdf) cut (Pagoda Arts, San Francisco, CA) from 1/8” black acrylic, which served as a light shield and a bearing surface for the hanging rhizotron. The other half of the polycarbonate sheets had 8 mm holes drilled in centered at the same location and were used as counterparts to the sheets with tops and hooks.

#### Boxes and holders

Similar to Rellán-Álvarez et al., 2015, rhizotrons were placed in black polyethylene boxes with acrylic holders during plant growth and imaging. Modifications were as follows: 12”W x 18”L x 12”H black polyethylene boxes (Plastic-Mart, Fort Worth, TX, Part number R121812A) were outfitted with a top holder and inner divider system to grow 12 rhizotrons at a time. The inside dividers were composed of 1/8” black acrylic cut into five sheets (bin-b.pdf, bin-c.pdf, and bin-d.pdf), which interlock perpendicularly with a large middle sheet (bin-d.pdf), as well as two smaller pieces (bin-b.pdf, bin-c.pdf) at each end. This divider system creates chambers, which isolate the rhizotrons from each other, thus preventing light affecting neighboring rhizotrons when one rhizotron is removed. Each bin has a bin top (rhiz-bin-top-enlarged.pdf), also cut from 1/8” black acrylic, has 12 cut outs arranged in two rows of six, where the rhizotrons sit. The cut outs have two vertical tabs, which were glued in using TAP Acrylic Cement (TAP plastics, CA) and provide placement guides for proper rhizotron insertion. The bin tops had two posts in diagonally opposite corners, which were connected with sticky copper tape (McMaster-Carr #76555A641) placed on the underside of each bin top to create a conductive path that the robotic arm used to find each bin and calculate the location of each rhizotron. Every post was made from an M5 stainless steel button head screw (McMaster-Carr #92095A484), 35 mm long - 13 mm OD 18-8 Stainless Steel Unthreaded Spacer for M5 screws (McMaster-Carr, #92871A276), 25 mm long - 10 mm OD 18-8 Stainless Steel Unthreaded Spacer for M5 screws (McMaster-Carr, #92871A093), and an M5 stainless steel lock nut (McMaster #93625A200). The screw went through the larger spacer, through the bin top, through the smaller spacer, and everything was held together by the nut. Each post extended down and into a 10 mm hole precisely positioned at the edge of each black box.

### Biological components

#### Six accessions

Bay, Col-0, Est, Sha, Tottarp, and Tsu-0 were transformed using the floral dip method (Zhang et al., 2006) with the *ProUBQ10::LUC2o* reporter, which consists of the UBQ10 promoter region and first intron amplified from Col-0 genomic DNA, the plant-codon-optimized LUC2 codon optimized coding sequence, and a plasma membrane-localized mCherry coding sequence driven by the 35S promoter (additional details in Rellán-Álvarez et al., 2015). Positive transformants were selected under a fluorescent dissecting scope (Leica M165 FC) using the mCherry marker visible in the mature seed. Bay, Col-0, Est, Sha, and Tsu-0 seeds were carried through to the T3 generation. Tottarp was used in the T2 generation.

#### Diversity panel

*Arabidopsis thaliana* accessions were ordered from the Arabidopsis Biological Resource Center (ABRC: CS78885) (Supplementary Table 4). For vernalization, plants were grown in pots filled with Pro-Mix PGX soil (Premier Tech, Canada) and placed in a growth cabinet with long day conditions (16-hr light, 8-hr dark) at 10 °C and 50% humidity using fluorescent bulbs with a light intensity of about 200 μEm-1s-1. Upon flowering, plants were transformed as described above and then moved to a 22 °C greenhouse with long day conditions and 150-250 μEm-1s-1 light intensity. Transformation of the 192 initial accessions yielded 187 positive T1 lines, 171 of which could be confirmed in the T2 generation. Ten positive transformants from each of the T2 lines were screened on plates for strong luminescent signal once primary root length reached approximately 1.5 cm, usually around two weeks, but up to 26 days. Seedlings with strong root signal were given a line-identification letter, then transferred to pots, and grown in the 10 °C growth cabinet until inflorescences emerged (27 to 109 days after transfer to soil depending on the accession), at which point all plants were photographed and moved to a 22 °C greenhouse until seeds could be harvested. Ultimately, 148 lines yielded at least 3 independent lines with strong root luminescence. Before starting an experiment, seeds were again screened for positive transformants using the mCherry seed coat marker.

### Growth method

#### Rhizotron preparation

Rhizotron preparation was done as described in Rellán-Álvarez et al., 2015, with minor modifications: a polycarbonate plastic sheet with 8 mm holes and a sheet with the black rhizotron top and metal hooks were each laid on the table. Since the acrylic rhizotron top extends from the top in every direction, those sheets were laid down on the building surface with the acrylic piece and hooks lying just-off the edge of the surface and the screws pointing up, which ensured that the sheet lay flat. To prevent excess buildup of soil, the spacers were inserted into the sheets with 8 mm holes, and “flags” (spacers modified with small pieces of material at the top to cover the bottom part of the hook and screw, Flags-final.eps) were inserted into the sheets with the attached tops. Using an electric paint sprayer (Wagner Spraytech Control Spray Double Duty HVLP Sprayer Model #0518050), a mist of water was applied to the sheets. Peat based Pro-Mix PGX soil (Premier Tech, Canada) was gently sifted over the sheets using a 2 mm sieve (US Standard Sieve Series N ° 10) and excess soil was gently shaken off. The mist, sift, shake procedure was done for a second time, which creates a two-layer thick surface of soil thin enough to see a small amount of light through when held up to a lightbulb or window. To prevent soil from falling out, a folded and trimmed piece of paper towel was moistened then placed at the bottom of each rhizotron. The spacers and flags were removed and the flags were replaced with clean spacers. The two sheets are carefully placed together and rubber U-channels were slipped on to each side. A small handful of sifted soil was placed into the groove in the rhizotron top and gently pushed down. Completed rhizotrons were hung in the boxes and, due to the increased size of the boxes, 3 liters of water were added to the bottom of each box.

#### Sowing and growth

Two transfer pipettes (each ∼2 mL) of Peters® Professional 20-20-20 General Purpose fertilizer were added to each rhizotron immediately after building. Fertilized rhizotrons in filled boxes were covered and left to soak overnight. Seeds were stratified for two days at 4 °C in distilled water. Three seeds were sown in the center of each rhizotron using a p200 pipette (Eppendorf) and a unique barcode was placed on the rubber U-channel. Rhizotrons were sprayed down with water and a clear acrylic sheet was placed on top of the box then sealed with packing tape to create a humid environment. Three days after sowing, clear sheets were unsealed and rhizotrons were watered with two transfer pipettes of water. The following day, the clear sheets were removed and rhizotrons were watered again. Rhizotrons were watered with two pipettes of water until nine days after sowing. Plants were thinned to one central plant five days after sowing. Plants were grown under long day conditions (16-hr light, 8-hr dark) at 22/18 °C (day/night) using LED lights (Valoya, C-series N12 spectrum) with a light intensity of about 130 μEm-1s-1. For the six accessions, plants were grown until 9 days after sowing (DAS) or 21 DAS and then were imaged every day until 31 DAS. Five replicates of each accession were grown for each treatment and plants were watered with luciferin every six days. For the diversity panel, the population was grown up, as a whole, six times, thus having temporal population replicates rather than internal replicates in order to reduce variation effects of growth condition. Accessions were always planted in the same locations relative to each other so the population was treated identically in each replicate. Plants were grown on shelves in the growth chamber next to the imager until 14 days after sowing (DAS). Subsequently, they were transferred into the imager and imaged every other day from 14 to 28 DAS with luciferin additions every six days.

#### Plant imaging

On each imaging day (timing depending on experiment), GLO-Bot loaded each rhizotron into the imager then closed the door and triggered μManager to capture 5 min exposures on each side of the rhizotron. Inside the imaging system, the rhizotrons were rotated using a Lambda 10-3 Optical Filter Changer (Sutter Instrument®, Novato, CA). Once root imaging was complete, a shoot image was taken using strips of LEDs and an ids UI-395xLE-C camera with a Fujinon C-Mount 8-80 mm Varifocal lens. If it was the first imaging day or a designated luciferin day (every six days), GLO-Bot added 50 mL of 300 μM D-luciferin (Biosynth International Inc., Itasca, IL) to the top of each rhizotron immediately before loading the rhizotron into the imager.

### Image analysis

Initial root image preparation to combine and align the four raw images was done as described in Rellán-Álvarez et al., 2015. Initial background noise removal was done using `remove.background.ijm`. By opening the correct blank image and then running this macro, the open blank image will be subtracted from all of the files in the folder being processed by the macro. The ImageJ plugin Template Matching and Slice Alignment was run as instructed in the manual to register all root images. Images are then run through the macro denoise_overlay.ijm to remove additional noise and add images together, which maximizes signal intensity throughout the root by: (1) subtracting 1.5 from all pixels to remove pixels with low values; (2) multiplying all values by 3, which greatly amplifies pixels with high values; (3) subtracting 8 from all values, which again removes pixels with low values (i.e. those that did not get amplified); (4) running the ImageJ command Despeckle, which replaces every pixel with the median value in its 3×3 neighborhood (Schindelin et al., 2012; Schneider et al., 2012); (5) setting and using a threshold to generate a binary image; (6) running the ImageJ commands Despeckle, Erode, and Dilate to further remove small pieces for noise; (7) using the ImageJ command Analyze Particles to get rid of small particles; and (8) merging the cleaned image with that of the previous day using the ImageJ command Add Create. This macro can be adjusted using the subtract-multiply-subtract sequence to change how signal intensity is amplified by manually testing the values on a handful of images.

Next, `macros invert.ijm` and `tip_tracking.ijm` are run to prepare images for GLO-RIAv2 (https://doi.org/10.5281/zenodo.5574925). This macro works by: (1) opening two consecutive images; (2) using the ImageJ command Dilate twice on the image from the earlier time point; (3) using the ImageJ command Subtract Create to subtract the earlier day from the later day. By dilating the earlier root system, we are able to ensure complete subtraction from the later time point and, therefore, only analyze the new root growth. Again, the thresholding in the new growth isolation can be refined if needed. Images were then run through GLO-RIAv2 and downstream analysis was performed in R (R Core Team, 2019) for additional cleanup and calculating traits. Initial formatting and combination of GLO-RIA output files is done using `1-format.Rmd`. This file uses the raw output, the “key”, which contains information about the plants, and the experimental start date to calculate relative imaging times (very important for comparing multiple experiments), extract identifying information for each rhizotron, and, for the local files, calculate the root angles with respect to gravity using trigonometric calculations based on whether the raw output is greater than or less than 90 °. Although we re-calculate the angles from 0 °-90 °, having the 0 °-180 ° angles tells us which way the vector is pointing and which trigonometric function should be used to calculate the end point for the root segment. It should be emphasized that angles are with respect to gravity and not with respect to the parent root. Formatted files are run through 2-clean.R to remove stray particles. This process works by (1) computing the distances between all root x-y points in an image; (2) using hclust(method = “single”) to cluster those distances; (3) split the cluster tree into two groups using cutree(); (4) calculating the distance between the groups and determining if it is greater or less than the predetermined proximity maximum; (5) if the distance is smaller than that distance the cluster is kept, if it is larger than the distance, cluster with the larger minimum distance to the center-x is kept and the other cluster is further examined; (6) in the cluster being examined, the cluster is again tested for whether the distance between the clusters is far too large (nearly twice that of the initial proximity maximum) or whether there are less than four points in the cluster, which would indicate the cluster is likely noise. If either of these are true, the cluster is discarded. If not, the cluster is further examined to test if the points are linear, which would indicate the cluster could be a root segment. The above sequence is done for one day, then the next day is added in and the process begins again, and this continues until only the “true root” segments remain. Parameter adjustments in this file primarily depend on the strength of the root signal and can be optimized through trial and error. The `2-clean.R` script outputs a text file describing which clusters were kept and removed, a pdf for each rhizotron showing all of the removal steps, two csv files, clean_removed_points.csv and clean_true_root.csv, which list all of the points that were either kept or removed, as well as two initial trait files: clean_ROIs.csv, and clean_traits.csv. Having these “true root” segments allows for calculation of traits at the new growth level as well as the whole root level. Some of these calculations are done during the root cleaning process, while others are re-calculated later in `3-summarize_traits.Rmd`.

Images were manually curated to eliminate those with very high levels of noise or roots that showed aberrant growth. Additionally, phosphorescent (“glow-in-the-dark”) stars in the upper right corner of some images were manually cropped out. Initially, these stars were placed in the images to aid image alignment, but instead generated too much noise.

### Trait processing

Raw trait measurements were converted into single values for each genotype in the software R (R Core Team, 2019) using the script `mcmcglmm_best.R`. This script is run for each trait, and the user inputs the raw phenotypic data for each trait. Several models were tested, including a second order polynomial for time, different error correction factors, and models with and without kinship-informed random factor. Each model was examined for parameter convergences using the Deviance Information Criteria (DIC), a Bayesian model version of Akaike Information Criterion (AIC). The selected model was chosen based on speed, simplicity and generalizability. While some traits might be better modeled using a polynomial function, we wanted to select a model that could be broadly applied to all traits over time rather than just one. The final model had the form:

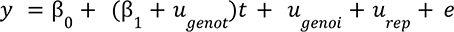

where the trait (*y*) and time (*t*) were fitted as fixed effects, genotype (u_genoi_), replicate (u_rep_), and the time and genotype interaction (u_genot_) were treated as random factors. The effects capture the average deviation for each genotype from the mean of the population. The script runs the data through a null model, which is a model run with no parameters and provides the baseline comparison for future models using the same input data, and then runs the previously described mixed model to estimate fixed effects by running 1,000,000 iterations in a Monte Carlo Markov Chain (MCMC) and 200,000 burn-in using the MCMCglmm R package (Hadfield, 2010). Since MCMCglmm utilizes a MCMC walk, the model has inherent stochasticity and can produce slightly different values each time it is run. For reproducibility, we set a “seed number” for the script then ran the model three times for each trait then selected the model with the best deviance information criteria (DIC) to calculate values fit for each genotype (i.e. breeding values) at timepoints 0, 48, 96, 144, 192, 240, 288, and 336 hours.

GEMMA (Zhou et al, 2014) was used for all Genome-Wide Associations (GWAs) of breeding values above. GEMMA was run as a linear mixed model (lmm) and with a kinship matrix to correct for accession relatedness. Only SNPs with a minimum allele frequency greater than 0.05 were examined. Bonferroni threshold, which was calculated using the number of linkage blocks within our population as computed by PLINK. Manhattan plots were generated in R.

## Notes

https://doi.org/10.5281/zenodo.5708422

https://doi.org/10.5281/zenodo.5574925

https://doi.org/10.5281/zenodo.5708430

https://doi.org/10.5281/zenodo.5709009

### Competing Interest Statement

The authors have declared no competing interest.

### Summary of Updates

Supplementary Data Table 3 has now been included. Acknowledgement of Wolfgang Busch has been added. Archived data DOIs have now been included.

